# Matrix mechanics governs mechano-metabolic adaptation across cancer grades in bladder spheroids

**DOI:** 10.64898/2025.12.03.692076

**Authors:** Sara Metwally, Dorota Gil, Justyna Śmiałek-Bartyzel, Joanna Pabijan, Gracjan Wątor, Kajangi Gnanachandran, Massimiliano Berard, Łukasz Kozłowski, Małgorzata Lekka

## Abstract

Extracellular matrix (ECM) mechanics critically influence cancer progression, yet the interplay between ECM viscoelasticity, architecture, and tumor cell adaptation remains incompletely understood. Here, we engineered collagen–hyaluronan hydrogels with tunable stiffness to mimic soft and stiff tumor microenvironments and studied bladder cancer spheroids representing benign, low-invasive, and highly invasive stages. Using hydraulic force spectroscopy, rheometry, and molecular analyses, we found that matrix stiffness differentially modulates spheroid morphology, migration, and expression of adhesion and metabolic markers. Active ECM remodeling via metalloproteinase MMP-2 facilitated migration in compliant but not rigid matrices, while mechano-metabolic coupling varied with cancer progression stage. These findings reveal how bladder cancer cells adapt to mechanical cues through coordinated biomechanical and metabolic responses, underscoring the importance of integrating cellular and matrix mechanics in modeling tumor invasion and developing targeted therapies.

## Introduction

The tumor microenvironment (TME) exerts a decisive influence on cancer initiation, invasion, and metastasis^1–3^. Among its constituents, the extracellular matrix (ECM) provides both structural and biochemical cues that shape tumor cell behavior. Collagen, the predominant ECM protein^4^, not only supports tissue integrity but also regulates adhesion, migration, and mechanotransduction^5,6^. Increasing evidence links ECM rigidity and architecture to tumor aggressiveness and resistance to therapy^7,8^, as elevated rigidity in matrices promotes invasion^9,10^, angiogenesis, and chemoresistance^11^, correlating with poor clinical outcomes^12^, and reduced drug efficacy^13,14^.

Cancer cells constantly interact with and remodel their surrounding ECM. Various studies have shown that metastatic cancer cells, together with stromal cells, regardless of their individual or collective migration mode^15,16^, interact with the ECM, frequently remodelling surrounding collagen matrix through deposition, degradation, and crosslinking of matrix components, which enhances local rigidity and facilitates tumor cell invasion^17,18^. These dynamic mechanical interactions generate diverse biomechanical cues—tension, compression, and shear—that are transmitted intracellularly through integrins and focal adhesions ^19,20^. Integrin activation triggers signaling cascades involving Rho GTPases and transforming growth factor-β (TGF-β), leading to cytoskeletal reorganization, gene regulation, and epithelial-to-mesenchymal transition (EMT). Mechanotransduction thus acts as a central mechanism linking ECM mechanics to tumor cell adaptation and invasiveness^21–23^.

Mechanical remodeling of the TME is closely intertwined with metabolic reprogramming, a hallmark of cancer progression^24,25^. To meet increased bioenergetic and biosynthetic demands, tumor cells often shift from oxidative phosphorylation to aerobic glycolysis (the Warburg effect), supporting proliferation and survival under stress^26,27^. Yet, despite mounting evidence connecting matrix stiffness to metabolic shifts, the reciprocal interplay between ECM viscoelasticity, structure, and tumor metabolism remains poorly defined. Viscoelasticity—a defining but underexplored property of the TME—modulates how cells perceive and respond to mechanical forces, influencing both tumor progression and therapeutic outcomes^28,29^. Understanding how these mechanical and rheological cues shape tumor behavior is essential for recreating physiologically relevant in vitro models and for advancing patient-specific drug testing platforms^30–32^.

Traditional two-dimensional (2D) cell cultures fail to capture the mechanical complexity of the TME, often leading to limited predictive value. Emerging three-dimensional (3D) models—such as spheroids, organoids, and tumor-on-a-chip systems—offer more realistic cellular interactions and gradients of oxygen, nutrients, and stiffness^33,34^. Among these, collagen–hyaluronic acid (Col–HA) hydrogels have emerged as versatile biomimetic platforms. Collagen provides a fibrillar scaffold and integrin-binding sites, whereas hyaluronan contributes hydration, viscoelasticity, and CD44- or RHAMM-mediated signaling^35,36^. Their combination enables tunable stiffness, porosity, and biochemical functionality, capturing essential features of native ECM mechanics.

Recent studies have demonstrated that Col–HA hydrogels modulate cancer cell invasion, angiogenesis, and matrix remodeling in a composition-dependent manner. For example, increasing HA content enhances proangiogenic factor expression and promotes invasive phenotypes, while variations in collagen density alter microarchitecture and migration modes^37,38^. Beyond structural mimicry, these matrices transmit mechanical and biochemical cues that can drive epithelial-to-mesenchymal transition (EMT^39^) and affect cell metabolism, making them powerful tools to probe tumor mechanobiology.

In this study, we engineered Col–HA hydrogels with tunable mechanical properties to emulate soft and stiff tumor microenvironments and investigated how these conditions regulate bladder cancer spheroid morphology, migration, and molecular signatures. Using hydraulic force spectroscopy (HSF) and rheometry, we quantified the viscoelastic behavior of spheroids derived from benign (HCV29), low-invasive (HT1376), and highly invasive (T24) bladder cell lines. We further analyzed key adhesion molecules, matrix metalloproteinases (here, MMP-2 and MMP-9), and metabolic enzymes to uncover the mechanisms underpinning cancer cell adaptation to different biomechanical contexts. Our results reveal that bladder cancer cells remodel their microenvironment via MMP-2–mediated degradation, enabling migration in compliant but not rigid matrices. We identify stiffness-dependent differences in adhesion and metabolic reprogramming that reflect progression-linked mechano-metabolic coupling. Together, these findings demonstrate how ECM mechanics regulate tumor growth and invasion through bidirectional feedback between cells and their matrix, providing mechanistic insight into the interplay between tumor biomechanics and metabolism.

### Morphological and mechanical differences in engineered collagen–hyaluronic acid hydrogels

Building on prior work^37^, Col–HA hydrogels fabricated from atelocollagen (pCol) and telocollagen (tCol) exhibited distinct structural and mechanical properties. Scanning electron microscopy (SEM) revealed that pCol–HA hydrogels possessed wider fibrillar spacing and fewer pores compared to tCol–HA (**Fig. 1A,B**). Quantitative analysis across 10 images showed 34 ± 16 pores versus 108 ± 59 (**Fig. 1C**), with corresponding median pore areas of 0.033 ± 0.020 µm² and 0.175 ± 0.081 µm² (**Fig. 1D**; median ± MAD). Despite heterogeneity, interfibrillar gaps remained below 2.5 µm, suggesting limited passive cell migration^40^ and a reliance on cell-mediated remodeling.

**Fig. 1.**
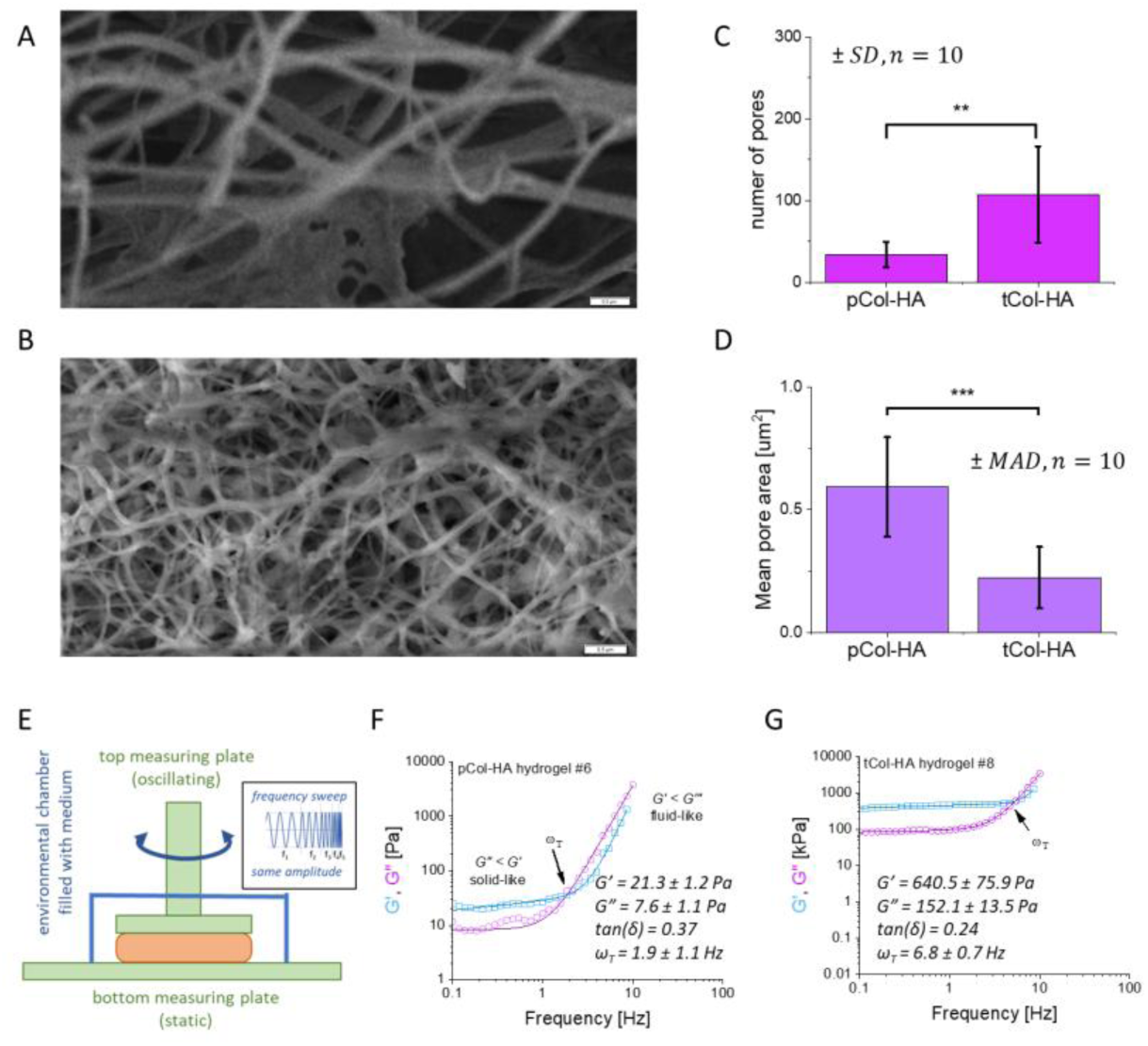
Morphological and rheological properties of pCol-HA and tCol-HA hydrogels. Representative SEM images of pCol-HA (A) and tCol-HA (B) reveal distinct pore architecture (scale bar = 0.5 µm). Quantification of pore density, i.e., the number of pores per image (C), and mean pore surface area (D) was performed on images (32µm² each). Data for pore density are shown as mean±SD (*n* denotes number of images); pore areas are presented as median±MAD (*n*=10). Statistical significance was determined by the Mann-Whitney nonparametric test (*α*=0.05; *\*\*p*<0.001, *\*\*\*p*<0.0001). Frequency-sweep rheology (E) yielded storage (*G′*) and loss (*G″*) moduli for pCol-HA (F) and tCol-HA (G) fitted with double-pawer law^44^ (exemplary hydrogels #6 and #8 are shown). The final rheological parameters: *G′, G″, ω_T_* (transition frequency between solid-like and fluid-like regimes), and loss tangent *tan(δ)=G″/G′* are calculated as a mean±SD (*n*=9 hydrogels per type).

Rheological profiling confirms substantial mechanical divergence (**Fig. 1E–G**; *n* = 9 hydrogels per group). pCol–HA hydrogels were markedly more compliant (storage modulus *G′* = 19.7 ± 9.1 Pa, loss modulus *G″* = 7.9 ± 2.9 Pa) in contrast to tCol–HA hydrogels (*G′* = 640.5 ± 75.9 Pa; *G″* = 152.1 ± 13.5 Pa). The higher loss tangent (*tan(δ) = G″/G′;* 0.37 vs 0.24, retained viscous components (20–30%) within the range characteristic for native soft tissues^41–43^) and lower transition frequency (*ω_T_* = 1.9 ± 0.2 Hz vs *ω_T_* = 6.8 ± 0.7 Hz) of pColHA, marking the changeover switch from solid- to fluid-like behavior, reflected a more compliant, loosely crosslinked matrix. Together, these results indicate that pCol–HA forms a softer, more remodelable microenvironment, whereas tCol–HA yields a stiffer network with denser, finer porosity.

### Bladder cancer spheroids display reduced rigidity and enhanced viscosity

Hydraulic force spectroscopy (HFS^45^, **Fig. 2A**) revealed distinct rheological signatures between malignant and non-malignant bladder spheroids. The storage modulus *G′* d followed a single power-law dependence (*G′(ω) = A_0_·ω^α^*, **Fig. 2B**). In contrast, the loss modulus *G″* exhibited a biphasic response, increasing at higher frequencies (above ∼8 Hz), with double power-law behavior 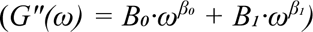. Fitted parameters are listed in the **Suppl. Table S2**. The prefactors *A_0_* and *B_0_* reflect low-frequency, soft-network elasticity, likely originating from a fractal or loosely crosslinked polymer network. Across frequencies, storage modulus ranked as *G′_HCV29_* > *G′_HT1376_* > *G′_T24_*, confirming that cancer spheroids are softer than non-malignant controls, consistent with previous AFM-based measurements on single cells and spheroids^46–48^. Consistently loss tangent taken from low-frequency linear region, was constantly higher for cancer spheroids: 0.45 ± 0.05 (HT1376) and 0.45 ± 0.12 (T24), compared to 0.38 ± 0.12 in HCV29 (the corresponding *p-*values i.e., *p* = 0.0016 and 0.0089, were obtained from unpaired Student t-test at the level of 0.05; **Fig. 2C**). Together, these data demonstrate that bladder cancer spheroids possess a softer, more viscous mechanical phenotype, consistent with enhanced deformability associated with malignant transformation.

**Fig. 2.**
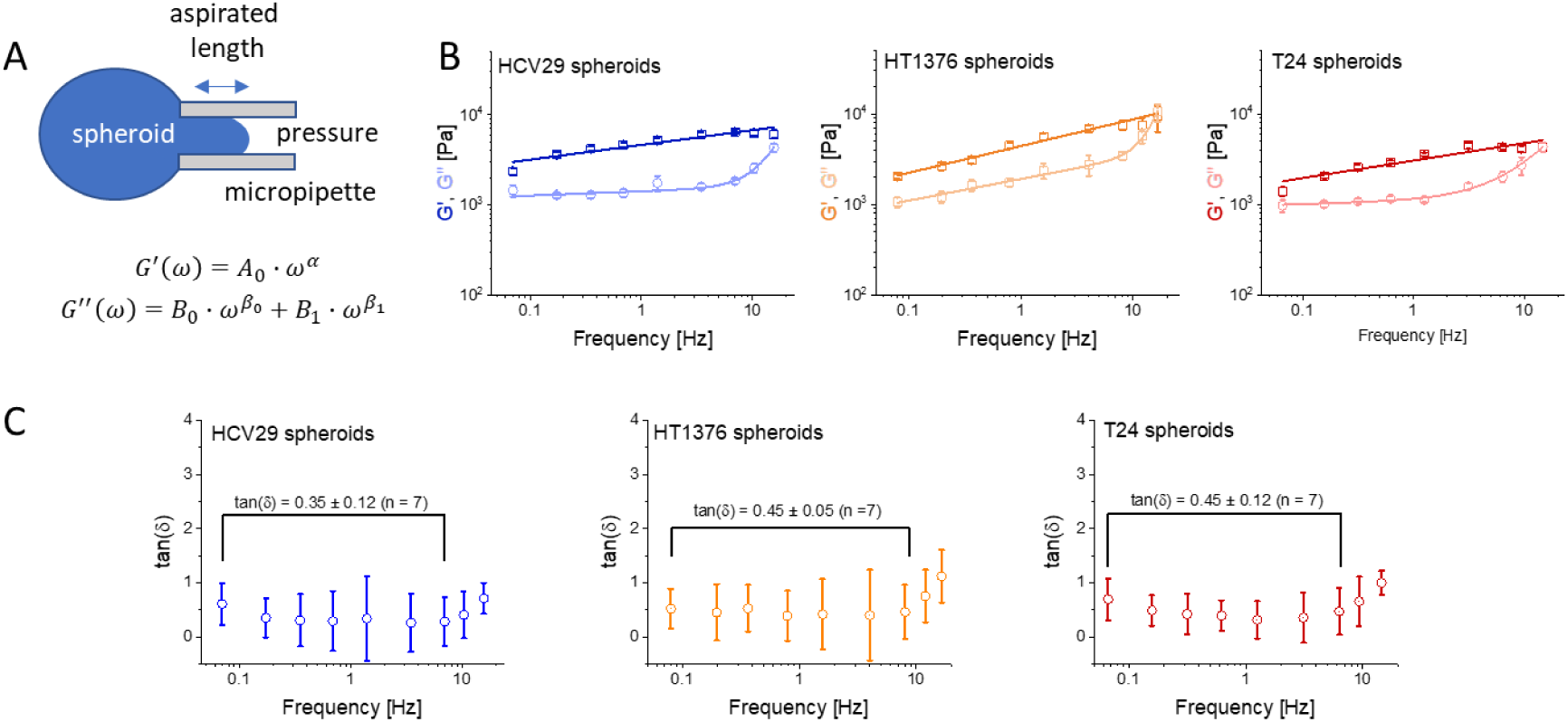
Rheological properties of bladder cancer spheroids and Col-HA hydrogels with embedded spheroids. (A) The idea of HFS-based rheological measurements alongside the applied power-law model. (B) Storage and loss moduli as a function of oscillation frequency (mean ± standard error (SE), n ≥ 10 spheroids). (C) An average loss tangent (*tan(α)*) calculated from the linear low-frequency region and represented as a mean value ± SD.

### Embedding spheroids does not alter initial hydrogel rheology

To assess whether incorporation of spheroids influences matrix mechanics, bladder spheroids were embedded within Col–HA hydrogels and cultured for 24h (cell viability inside spheroids is presented in **Suppl. Fig. S1**). Each hydrogel (*Ø* = 8 mm, thickness = 2 mm, volume = 100 mm³) contained 48 spheroids, with a density of ∼1–2 spheroids/mm², verified via optical imaging. This strategy avoided premature cell migration, commonly observed when spheroids form *in situ*^49,50^. Frequency sweep rheology (0.1–10 Hz; **Suppl. Note S2; Suppl. Fig. S2**) revealed no significant differences in storage modulus, loss modulus, or loss tangent compared to cell-free controls, indicating that initial viscoelastic properties remained unchanged (**Suppl. Fig. S2B**). Transition frequencies between solid-like and fluid-like regimes were likewise unaffected (pCol-HA: 2.0 ± 0.2 Hz vs. 1.9 ± 0.2 Hz; tCol-HA (6.8 ± 0.7 Hz in both control and spheroid-laden tCol–HA hydrogels, consistent with prior reports^51^). These results confirm that the intrinsic rheological behavior of both hydrogel formulations is preserved upon spheroid embedding.

### Rheological mismatch shapes spheroid morphology and energetic requirements

Subsequent analyses, therefore, focused on how these mechanically distinct matrices influence spheroid morphology, assessed via quantitative image-based shape analysis of elliptical spheroids and their contiguous migratory zones. To this end, phase-contrast microscopy images were analyzed using a custom image processing pipeline. Given that all spheroids exhibited an approximately elliptical shape, each was fitted with an ellipse (*n* = 25; **Fig. 3A**), including a zone of migrating cells that maintained structural continuity with the parent spheroid. The morphological parameters were extracted (**Suppl. Table S1**).

**Fig. 3.**
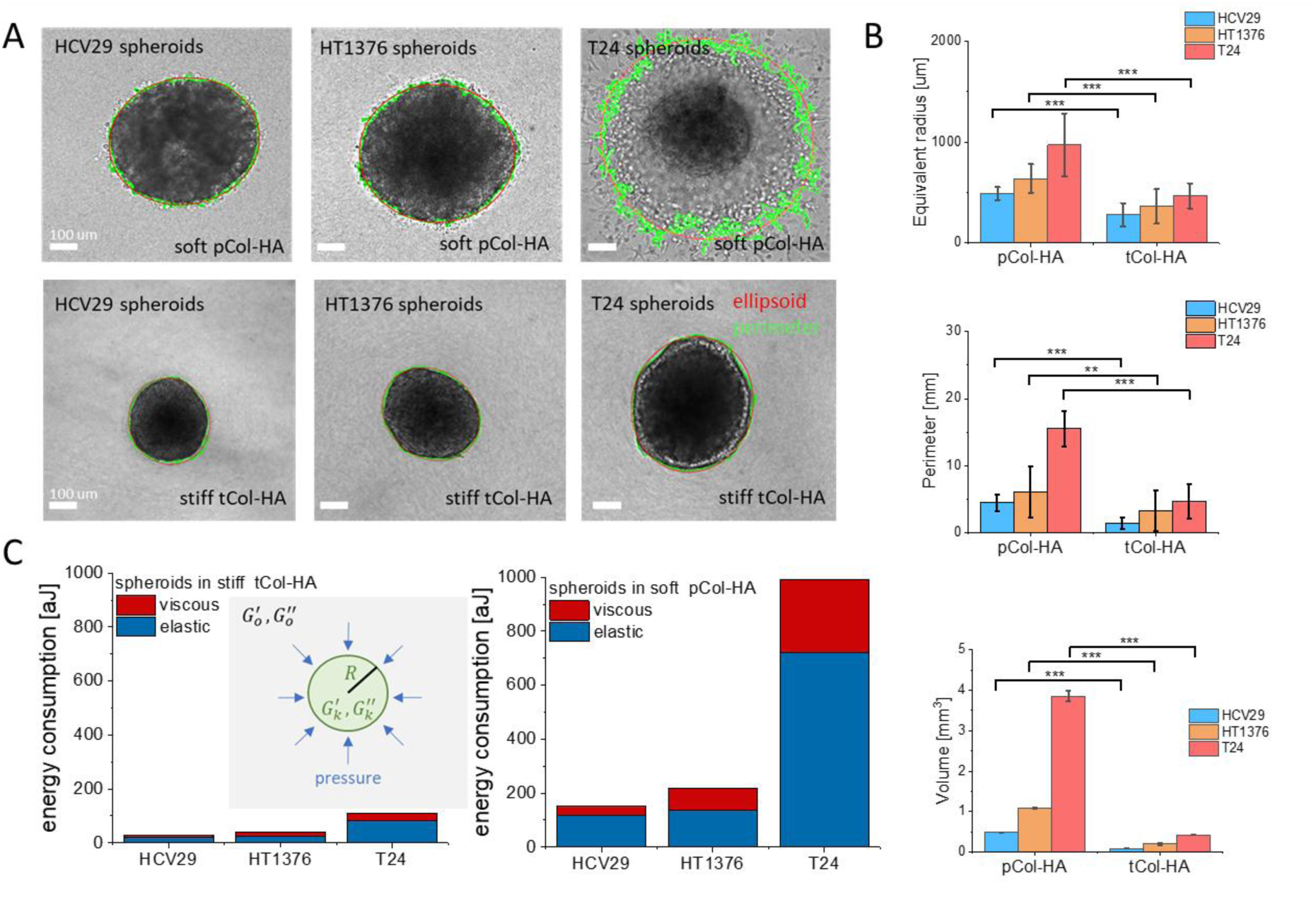
Morphology of bladder cancer spheroids in soft and stiff Col-HA hydrogels. (A) Fitting ellipsoids (red line) to each spheroid present in the optical images (scale bar 100 µm), quantified spheroid morphology together with a zone of migrating cells that maintained structural continuity with the parent spheroid, acquired after 24h of culture within Col-HA hydrogels. (B) Differences in perimeter and equivalent radius (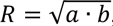, where *a* and *b* are major and minor axes), which was used to calculate spheroid volume (*n* = 25 spheroids). Statistical significance was verified using an unpaired t-test (*α* = 0.05), *\*\*p*<0.001, *\*\*\*p*<0.0001. (C) Theoretical model applied to estimate the energy required to maintain the spheroid shape in soft and stiff hydrogels, assuming 1% radial spheroid expansion over 1h.

Despite comparable aspect ratios (1.075 to 1.179), spheroid morphology diverged markedly across hydrogel conditions in a cell line–dependent manner (**Fig. 3B**). In soft pCol–HA hydrogels, spheroids displayed increased perimeter and volume, following the trend: *HCV29 < HT1376 < T24*. While HCV29 and HT1376 spheroids primarily expanded in volume, T24 spheroids exhibited extensive cellular migration into the surrounding matrix after 24h (**Fig. 3A**), indicating that soft hydrogels facilitate both expansion and invasion. Conversely, stiff tCol–HA hydrogels constrained spheroid growth, yielding smaller spheroid volumes and perimeters, consistent with mechanical confinement by the dense network.

To quantify the energetic cost of maintaining spheroid structure under these conditions, we developed a simplified model of a viscoelastic sphere embedded within a viscoelastic matrix (see **Suppl. Note S3**). This model accounts for both elastic and viscous contributions used to estimate the energetic cost of maintaining spheroid structure. Under steady-state conditions, only elastic energy is involved; during growth, viscous dissipation arises due to matrix resistance to expansion. Assuming a 1% radial expansion over 1 hour, we estimated the energy required to maintain spheroid shape (**Fig. 3C**). The model predicted greater energetic demand in the softer pCol–HA hydrogels than in the stiffer tCol–HA matrices. This counterintuitive result suggests that the energetic cost of spheroid maintenance arises not simply from matrix stiffness but from rheological mismatch between the spheroid and its surrounding microenvironment.

### Bladder cancer cell migration depends on matrix stiffness and collagen remodeling

We next examined how collagen network architecture governs bladder cancer cell migration. We compared cell motility on HCV29 monolayers and within soft and stiff 3D Col–HA hydrogels (**Fig. 4A**).

**Fig. 4.**
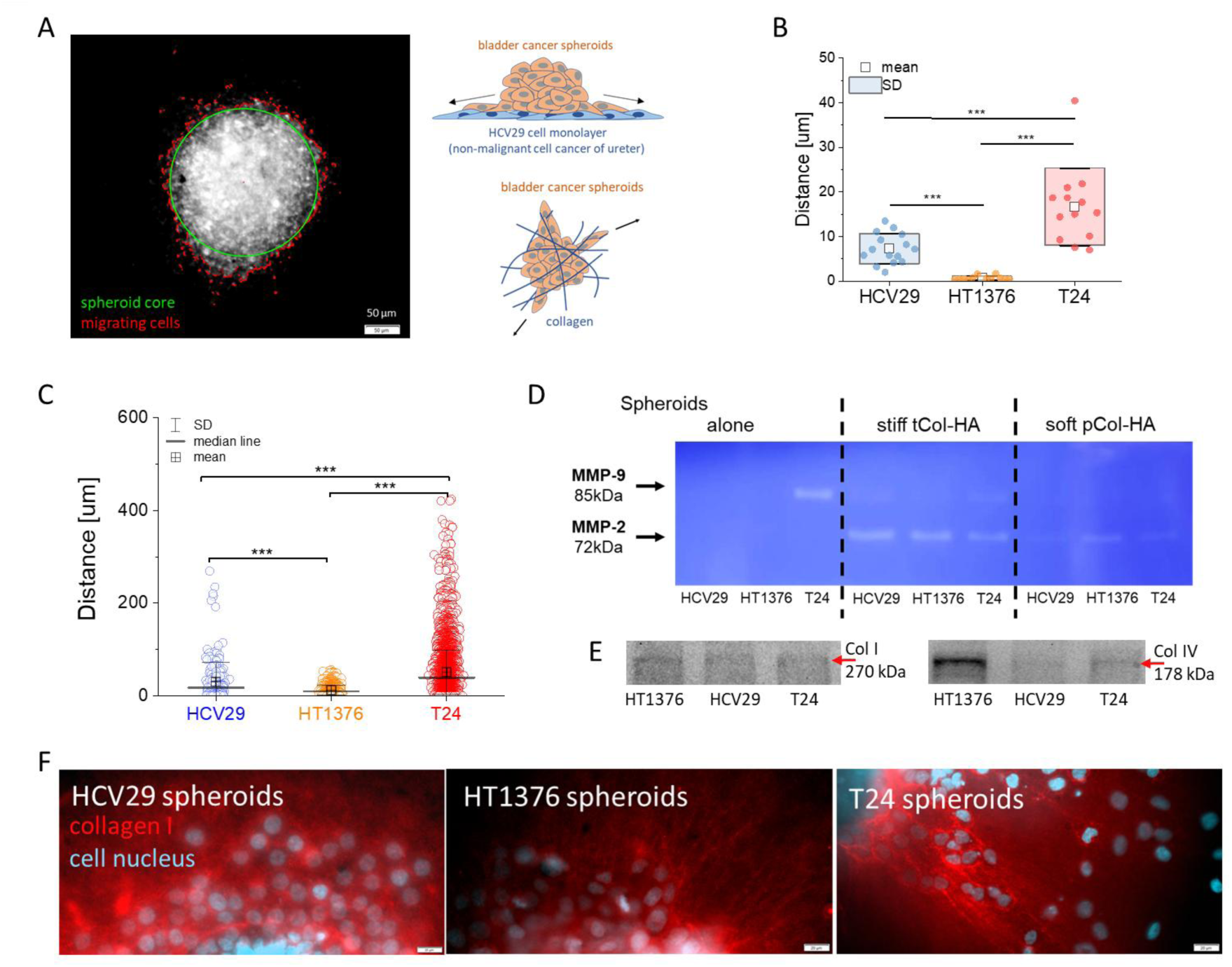
Cell escape from bladder cancer spheroids in Col–HA hydrogels of varying stiffness. (A) Quantification of cell migration, measured as the distance from the spheroid edge for cells migrating from (B) bladder cancer spheroids seeded on a monolayer of non-malignant HCV29 cells, and from spheroids embedded within soft 3D Col-HA hydrogels. Statistical significance in (B&C) was evaluated using the Kruskal-Wallis ANOVA test at α = 0.05; *\*\*\*p*<0.0001). (D) MMP-2 and MMP-9 activity in supernatants collected after 24h culture of spheroids in stiff or soft Col–HA hydrogels. (E) Collagens (Col I and Col IV) expression in bladder cancer cell lines. (F) Collagen matrix remodeled by spheroids in soft Col–HA hydrogels (nuclei, blue; collagen fibers, red; scale bar, 20µm).

Across all environments, cells exhibited a consistent migration hierarchy (T24 > HCV29 > HT1376, **Fig. 4B&C**) that persisted over 72 h, despite cell-to-cell variability (**Suppl. Fig. S3 & S4**). Given that hydrogel pore sizes fall below the limit for passive migration, movement was inferred to rely on active ECM remodeling. We therefore examined the activity of matrix metalloproteinases (MMP-2 and MMP-9), enzymes that primarily degrade denatured collagens and type IV collagen of basement membranes, but can also contribute, albeit less efficiently, to collagenI breakdown^52,53^. Zymographic analysis revealed distinct metalloproteinase activity profiles of MMP-2 and MMP-9. They were undetectable in HT1376 and HCV29 spheroids, while T24 spheroids expressed MMP-9 (**Fig.4D**). In spheroid supernatants, originating from spheroid cultures in soft and stiff Col–HA hydrogels, MMP-2 predominated, with activity enhanced in stiffer matrices, indicating a stiffness-dependent remodeling response. Collagen degradation by these enzymes permitted cell translocation in soft matrices but not in stiff hydrogels, where cells remained confined even after extended culture (**Suppl.Fig.S3&S4**). Immunoblotting confirmed that spheroids also synthesized collagens I and IV, supporting their direct role in matrix remodeling (**Fig. 4E**). Thus, we fluorescently stained collagen fibers around spheroids (**Fig.4F**). Fluorescent imaging of collagen fibers revealed a disordered network surrounding HCV29 spheroids, in contrast to the radially aligned, star-like organization induced by cancer spheroids. Migration correlated not with the formation of collagen tracts but with localized pericellular collagen deposition, most pronounced in HCV29 and T24 cells. Collectively, these findings indicate that bladder cancer cell migration in 3D hydrogels is jointly governed by matrix stiffness and the capacity for enzymatic and structural ECM remodeling mediated by MMP-2 and MMP-9.

### Matrix stiffness and density shape adhesion and metabolic marker expression

In the final phase of our study, we examined selected adhesion- and metabolism-related markers at the protein and mRNA levels. Immunoblotting focused on vinculin being a focal adhesion marker54, responsible for cell attachments to the surrounding environment, and on E- and Ncadherins (we already showed the presence of these cadherins in our earlier studied cells55), mediators of cell–cell adhesion that maintain spheroid cohesion^56^. Transcript analysis targeted HK2 (metabolic reprogramming marker^57,58^), and SDC4 (overexpressed in bladder cancer cells^59^). Expression profiles revealed spheroid–type–dependent regulation of these markers, reflecting differential engagement of adhesion and metabolic pathways across bladder cell lines (**Fig. 5**). Detailed quantifications for control and hydrogel-embedded spheroids are provided in **Suppl. Fig. S5&S6**).

**Fig. 5.**
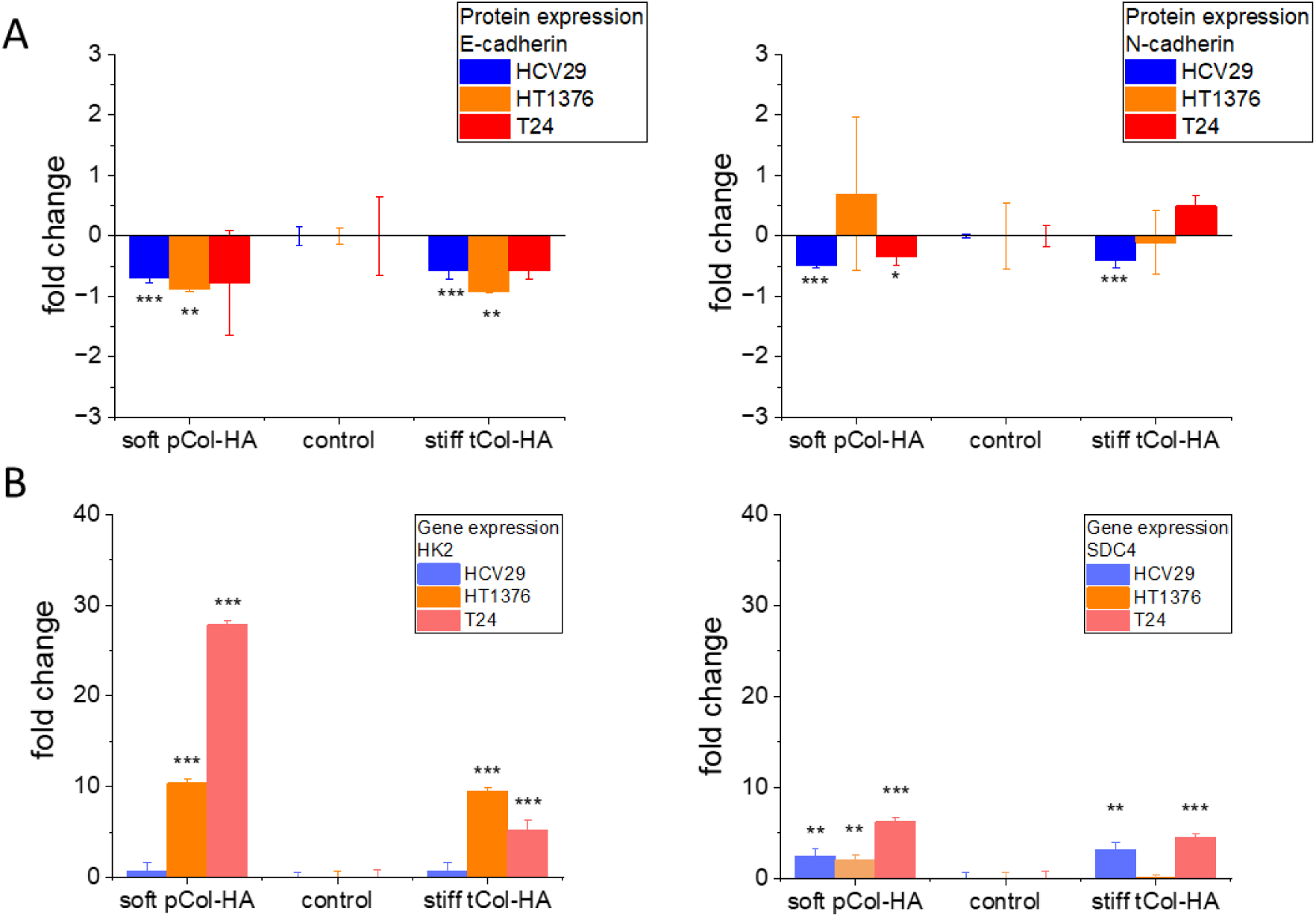
Expression profiles of specific adhesion- and metabolism-related markers in bladder cancer spheroids. Spheroids representing three bladder cancer cell lines (HCV29, T24, HT1376) were embedded in collagen–hyaluronan (Col–HA) hydrogels. Relative expressions of E-cadherin, N-cadherin, hexokinase 2 (HK2), and syndecan-4 (SDC4) were normalized to control (i.e., expressions in spheroids grown only in culture medium, *n* = 3) and converted to fold change. Statistical significance was obtained by unpaired Student’s t-test at the level of 0.05 (\**p < 0.01, **p*<0.001, *\*\*\*p*<0.0001).

Vinculin, a focal adhesion protein, was analyzed to assess mechanosensitive responses within compliant and rigid Col–HA matrices. In non-malignant HCV29 spheroids, vinculin expression increased in stiff hydrogels (**Suppl. Fig. 4**), suggesting enhanced anchoring to the denser collagen network. In contrast, vinculin levels in cancerous HT1376 and T24 spheroids remained unchanged across stiffness conditions (**Suppl. Fig. S4**), indicating that focal adhesion reinforcement is not a dominant mechanoadaptive mechanism in bladder cancer spheroids embedded in 3D matrices. Cadherin profiling revealed subtype-specific expression consistent with prior observations^55^: HCV29 cells expressed both E- and N-cadherin, T24 cells predominantly N-cadherin, and HT1376 cells exclusively E-cadherin. Both adhesion molecules contribute to spheroid cohesion^60,61^, yet their expression was variably modulated by matrix mechanics. E-cadherin levels were markedly reduced in HCV29 and HT1376 spheroids (**Fig. 5A**), regardless of the matrix rheology, while T24 spheroids – characterized by minimal baseline E-cadherin – showed no significant changes. N-cadherin expression decreased in HCV29 spheroids across both hydrogels, was undetectable in HT1376 (due to minimal initial expression), and exhibited a stiffness-dependent shift in T24 spheroids, decreasing in soft and increasing in stiff matrices (**Fig. 5A**). Consistent with the presence of extracellular matrix components between cells, spheroids secreted collagen (**Fig. 4E**) and fibronectin^62,63^, the latter engaging integrins via syndecan-4, which is expressed by all examined bladder cancer lines. Together, these findings indicate that spheroid structural integrity and adhesion dynamics arise from coordinated modulation of cadherin expression and ECM–integrin–syndecan interactions within stiffness-defined microenvironments. To probe metabolic activity, we analyzed HK2 expression, encoding hexokinase-2—the enzyme catalyzing the first and rate-limiting step of glycolysis. In line with its established oncogenic upregulation^64^, HK2 levels remained stable in non-malignant HCV29 spheroids but were markedly elevated in HT1376 and T24 spheroids embedded in both compliant and stiff Col–HA hydrogels (**Fig. 5B**). Given that syndecan-4 (SDC4) is overexpressed in bladder cancer^59^ and participates in metabolic reprogramming via AKT/mTOR and mechanotransduction pathways^65^, we next assessed SDC4 expression. We observed a pronounced upregulation in HCV29 spheroids within soft and stiff Col-HA, in HT1376 spheroids within soft Col-HA, and in T24 spheroids under both compliant and rigid conditions (**Fig. 5B**), highlighting a stiffness- and subtype-dependent regulatory pattern.

Together, these findings reveal distinct metabolic and adhesion signatures across spheroid types. Non-malignant HCV29 spheroids modulate vinculin and SDC4 expression in response to matrix mechanics, whereas cancerous spheroids display stable vinculin levels but upregulated HK2 and SDC4, consistent with enhanced glycolytic activity and mechanometabolic adaptation within 3D microenvironments.

## Discussion

In this study, we designed collagen–hyaluronic acid (Col–HA) hydrogels with tunable mechanical properties to model soft and stiff tumor-like niches and investigate their effects on bladder cancer spheroid morphology, migration, and molecular responses. Spheroids derived from three bladder cell lines lines (**Suppl. Note S3)** exhibited distinct behaviors determined by both their intrinsic characteristics and the rheological properties of the surrounding matrix— mirroring progressive stages of bladder cancer development. Specifically, non-malignant ureter-derived HCV29 cells and two bladder cancer lines—HT1376 (carcinoma) and T24 (transitional cell carcinoma)—served as models of low- and high-grade malignancy, respectively. This system thus enabled controlled exploration of how microenvironmental stiffness modulates spheroid mechanics, invasiveness, and mechano-metabolic adaptation across bladder cancer progression.

The three bladder cell lines displayed distinct biomechanical and molecular responses to soft and stiff Col–HA hydrogels, reflecting their intrinsic rigidity and invasiveness. Nonmalignant HCV29 spheroids formed the most rigid and compact structures. In soft matrices, they expanded moderately due to reduced confinement but showed no change in HK2 expression, indicating minimal glycolytic reprogramming^66^. Their migration was limited— greater than HT1376 but less than T24—and driven primarily by adhesion remodeling rather than metabolic activation or proteolysis, as evidenced by decreased cadherin and increased SDC4 expression. The absence of MMP-2 activity and randomly oriented collagen fibers suggested that HCV29 spheroids rely on intrinsic rigidity and cell–matrix adhesion to maintain structure. In stiff hydrogels, external mechanical support and insufficient proteolytic activity reduced energetic demand, stabilizing spheroid shape but further limiting motility. Carcinoma HT1376 spheroids also remained compact but exhibited higher HK2 expression and MMP-2 activity in soft matrices, consistent with increased energy demand and attempted ECM remodeling. Despite these molecular changes—together with reduced cadherins and elevated SDC4—their migration remained minimal. Collagen fibers arranged radially from the spheroid indicated partial remodeling, yet not enough to enable invasion. In stiff matrices, HT1376 spheroids were non-migratory; while MMP-2 levels were elevated, the dense collagen network resisted degradation. The energy cost of maintaining spheroid integrity decreased with stiffness, but metabolic activation and adhesion remodeling were insufficient to overcome mechanical constraints, indicating a decoupling between invasive priming and actual motility. In contrast, transitional cell carcinoma T24 spheroids—the softest and most deformable—displayed robust migration in soft hydrogels. They integrated high glycolytic activity (HK2 upregulation), strong SDC4 induction, and moderate MMP-2 activity to coordinate ECM remodeling and movement. Migrating cells were surrounded by reorganized collagen sheaths, reflecting active deposition and fiber alignment. Low E-cadherin and high N-cadherin expression indicated weakened intercellular cohesion supporting motility. In stiff matrices, migration was suppressed despite elevated MMP-2 and SDC4 levels, suggesting active but ineffective attempts at remodeling under mechanical constraint. The increased energetic cost of maintaining spheroid structure and altered cadherin balance (E-cadherin↓, N-cadherin↑) pointed to collective stabilization rather than individual migration. Overall, these results demonstrate that bladder spheroid behavior emerges from an interplay between intrinsic mechanical properties, matrix stiffness, and molecular adaptation. HCV29 spheroids rely on rigidity and adhesion, HT1376 spheroids activate metabolic and proteolytic pathways without achieving invasion, and T24 spheroids effectively couple metabolic reprogramming, adhesion remodeling, and ECM interaction to drive migration in compliant but not rigid microenvironments.

Based on the obtained data, we focus on elaboration of the mechanism, explaining the migration of cells from bladder cancer spheroids (**Fig. 6**).

**Fig. 6.**
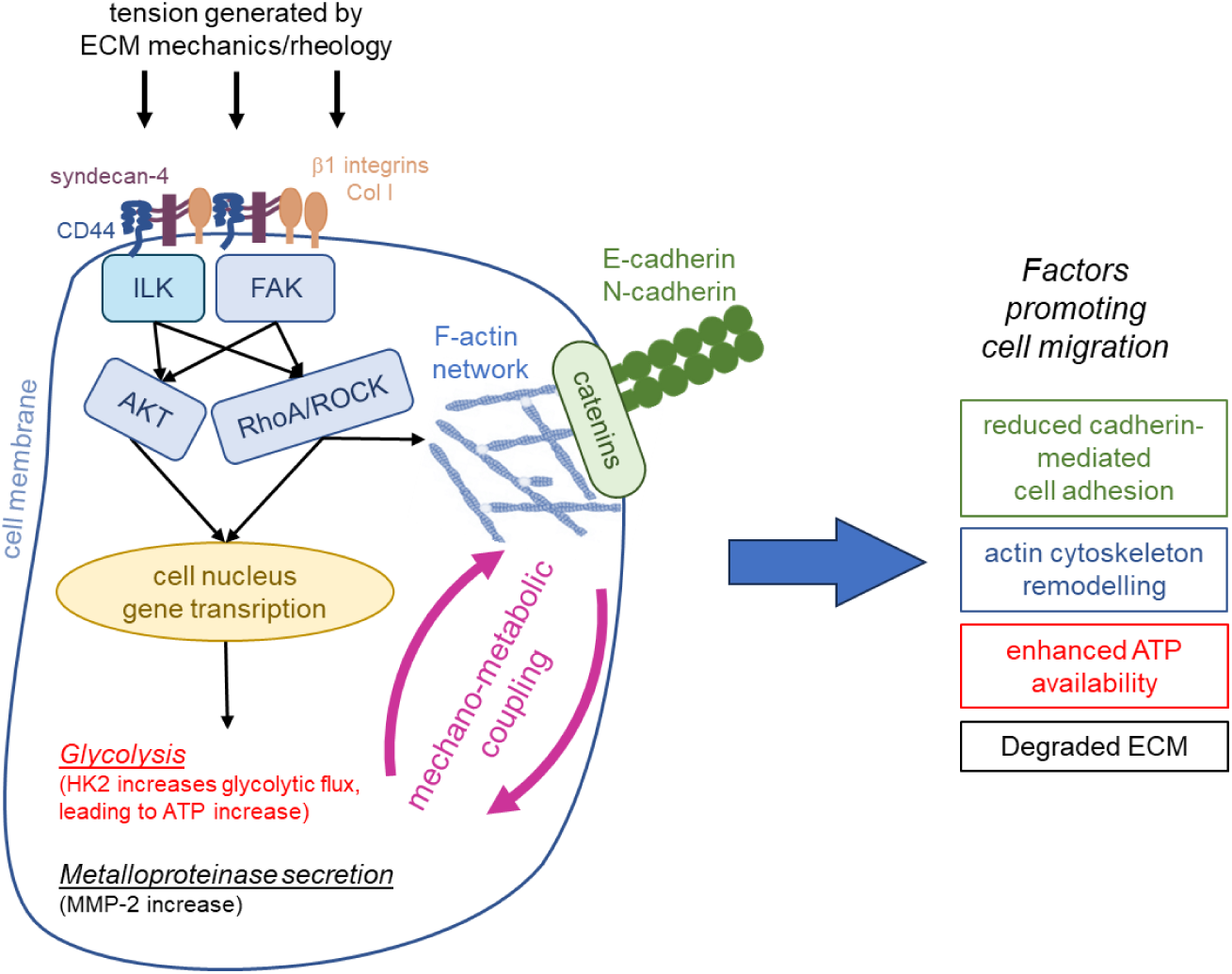
Schematic representation of the interplay between extracellular matrix (ECM) mechanics and cell metabolism. Mechanical tension from ECM rheology activates integrins and syndecans, initiating FAK/ILK-mediated signalling through AKT and RhoA/ROCK pathways. These cascades remodel the actin cytoskeleton, regulate E- and N-cadherin-dependent adhesion, and induce gene programs such as MMP-2 upregulation to promote matrix ECM degradation and cell migration. Simultaneously, mechanotransduction enhances HK2 expression, boosting glycolysis and ATP level to support cytoskeletal remodeling and cell motility.

The 3D Col–HA microenvironment exerts mechanical forces transmitted from the spheroid surface into its core, generating tension on mechanosensitive surface receptors such as integrins and syndecans. Building on our previous findings that SDC4 is overexpressed in bladder cancer cells compared with non-malignant HCV29 cells^59^, we now show that SDC4 expression increases further when spheroids are embedded in soft hydrogels. This upregulation likely activates an SDC4/integrin→FAK→RhoA/ROCK signaling cascade, promoting cytoskeletal remodeling in response to local mechanical stimuli^73^. Such mechanotransduction leads to the destabilization of adherens junctions and downregulation of cadherins^74^—consistent with our observations—via transcriptional repression and post-translational mechanisms. Concurrently, radial alignment of collagen fibers facilitates directional migration, while coordinated modulation of SDC4, cadherins, and HK2 expression enables mechanical and metabolic adaptation across matrix environments. Elevated HK2 supports enhanced glycolytic flux, providing energy for both motility and matrix remodeling, which is reflected by increased MMP-2 activity and collagen degradation. Our findings highlight a mechanistic link between matrix stiffness, SDC4-mediated mechanotransduction, and metabolic reprogramming that underlies bladder cancer spheroid plasticity and invasive potential in 3D microenvironments.

## Acknowledgements

This work was supported by the National Science Center (Poland), project NCN-OPUS no UMO-2021/41/B/ST5/03032. SEM imaging (Fig. 1A&B) and the protocol for staining spheroids embedded in Col-HA hydrogels (Fig. 4G) were developed within the project NCNMiniatura 7 no DEC-2023/07/ X/ST5/00688.

## Authors contributions

SM, JP, JSB, and KG conducted sample preparations; DG and JP performed Western blots; GD conducted zymography for metalloproteinase; SM carried out SEM, rheometer measurements; KG and MB conducted the HFS measurements; MB performed HFS data analysis; JP conducted experiments of spheroids on cell monolayer, SM and JP contributed to fluorescent imaging and staining; GW conducted mRNA isolation and gene expression experiment and analysis, LK wrote MATLAB code for migration analysis; ML wrote designed the project, analyzed data, wrote Python code for spheroid morphometry and energy demand analysis, and wrote the initial manuscript draft. All authors edited, revised, and approved the manuscript.

## Code availability

Codes supporting the findings of this paper are available from the corresponding author upon reasonable request.

## Materials and Methods

### Cell cultures

Three human bladder cancer cell lines were used in the study, namely, HCV29 – non-malignant cancer cells of the ureter (established from the Institute of Experimental Therapy, PAN, Wroclaw, Poland), T24 – transitional cell carcinoma, and HT1376 – grade III bladder carcinoma (ATCC, LGC Standards, Poland). Cells were cultured in tissue culture flasks (25 cm^2^ surface area, TPP, Genos, Poland). Roswell Park Memorial Institute 1640 culture medium (RPMI, Merck, Poland) supplemented with 10% Fetal Bovine Serum (FBS, ATCC, USA) was used for HCV29 and T24 cells, and Eagle’s Minimum Essential Medium (EMEM, ATCC, USA) with 10% FBS for HT1376 cells. Subculturing was performed every 3–4 days when the cell monolayer reached full confluence. Cells were centrifuged (1800 rpm, 4 min) after trypsinization with 0.25% trypsin–EDTA solution (Merck, Poland) in phosphate-buffered saline (PBS, Merck, Poland). Next, the supernatant was removed, and the pellet was resuspended in fresh culture medium (with 10% FBS) and seeded into new culture flasks. Optimal culture conditions, i.e., 37 °C, 5% CO_2_, and 95% humidity, were provided in a CO_2_ incubator (Nuaire, USA).

### Spheroid growth

Bladder cancer spheroids were formed in 96-well U-bottom 3D cell culture plates (ThermoFisher Scientific, USA) by seeding cells at densities of 5000 cells/well (HCV29), 9000 cells/well (T24), and 9000 cells/well (HT1376), counted using a Burker chamber (Bionovo, Poland). The density of cells was adjusted to obtain spheroids with a final diameter between 350 and 400 µm. To grow spheroids, 150 µL of the cell suspension was added to each well filled with the cell-type corresponding culture media, i.e., RPMI 1640 (HCV29, T24 cells) and EMEM (HT1376 cells).

### Fabrication of hydrogels composed of collagen and hyaluronic acid (Col-HA hydrogels)

The preparation of Col-HA hydrogels follows the procedure published elsewhere^1^. Briefly, collagen hydrogels (collagen type I from bovine; 3 mg/mL PureCol (pCol) and 10 mg/mL TeloCol (tCol); Advanced BioMatrix, Cell Systems, Germany) were prepared by mixing cold (∼ 4 °C) collagen solution with ten-fold concentrated phosphate-buffered saline (PBS, Merck, Poland) in a volumetric ratio of 8:1. The pH of such solution was adjusted to 7.2 – 7.5 using 115 µL of 0.1 M NaOH solution (Avantor Performance Materials, Poland) dissolved in deionized water (dH_2_O, Direct-Q 3 UV, Merck, Poland). The water was filtered using a syringe filter with a pore size of 0.22 µm (TPP, Genos, Poland) to obtain sterile dH_2_O, which was then added to the solution to obtain a final volumetric ratio of 8:1:1 for collagen, PBS, and water. Collagen-hyaluronic acid hydrogels (i.e., Col-HA hydrogels) were prepared by diluting 10 mg of hyaluronic acid (HA, Advanced BioMatrix, Cell Systems, Germany) with 1 mL of ten-fold concentrated PBS. Collagen solution (pCol or tCol) was then mixed with the HA solution in an 8:1 v/v ratio. The pH of the solution was adjusted to 7.2–7.5 using 115 µL of 0.1 M NaOH, followed by the addition of sterile dH_2_O to obtain a final solution of collagen, hyaluronic acid, and water in an 8:1:1 v/v ratio. For Col-HA hydrogels, final concentrations are 2.4 mg/mL (for pCol) or 8 mg/mL (for tCol), and 1 mg/mL (for HA). All steps of Col-HA hydrogel preparation were conducted on ice to prevent gelation during the preparation process. Finally, 1 mL of Col-HA hydrogel solution was transferred into each well of the 24-well plates (TPP, Genos, Poland), which were placed in the CO_2_ incubator (Nuaire, USA), providing an atmosphere composed of 5% CO_2_ and 37°C for 120 min (pCol) or 60 min (tCol). Then, the hydrogels were incubated in a culture medium corresponding to the cell type used at 37 °C for 24h in a CO_2_ incubator (Nuaire, USA).

### Incorporation of spheroids into 3D Col-HA hydrogels

After 3 days of spheroid culture, they were collected from the U-bottom plates, and the specific numbers of spheroids were placed on top of 1 mL of an unpolymerized hydrogel solution in individual wells of 24-well plates. After polymerization (37 °C, 5% CO_2_), the prepared samples were incubated for 24 h in the culture medium corresponding to the cell type used. The number of incorporated spheroids was dependent on the measurement type.

### Extraction of spheroids from Col-HA hydrogels for qPCR and Western blot

For Western Blot and quantitative Polymerase Chain Reaction (qPCR), spheroids were isolated from Col-HA hydrogels. First, 96 spheroids of HCV29 and T24 cells, and 288 of HT1376 cells, were embedded in individual Col-HA hydrogels and incubated (37°C, 5% CO_2_) for 24h in culture medium corresponding to the cell type used. The number of spheroids was adjusted to obtain similar protein levels for all spheroid types in the blots. Next, a collagenase solution was prepared by dissolving collagenase type I (Life Technologies, USA) in Hanks’ Balanced Salt Solution with Ca^2+^ and Mg^2+^ (HBSS, Gibco, USA) in a 1:100 v/v ratio to obtain a concentration of 10 mg/mL (enzymatic activity 1U/µL). Next, the supernatant from the spheroid culture was removed, followed by rinsing in PBS. Hydrogels with spheroids were transferred into 2 mL Eppendorf tubes, and 1 mL of collagenase solution was added to each sample. Aftewards, they incubated for 90 minutes at 37°C, until the Col-HA hydrogel was completely dissolved. After adding 1% FBS, spheroids were centrifuged (1800 rpm, 4 min), moved to a fresh HBSS solution supplemented with 1% FBS, and centrifuged again. Next, the supernatant was removed, and all spheroid pellets were frozen at -80°C.

### SEM imaging of Col-HA hydrogels

The 1 mL of Col-HA hydrogel solution was polymerized in 24-well plates as described above, and incubated with PBS for 24 h. Next, PBS was removed, and the samples were fixed with a 0.5% glutaraldehyde aqueous solution (Avantor Performance Materials, Poland), followed by three rinses in dH_2_O. Next, the hydrogels were freeze-dried by rapid immersion in liquid nitrogen and lyophilized in the chamber under a controlled pressure of 10^−5^ Pa for 12 hours. Dehydrated samples were mounted on holders specific for scanning electron microscope (SEM), using double-sided carbon tape and coated with a 5 nm gold layer using a rotary-pumped sputter coater (Q150RS, Quorum Technologies, UK). To visualize the hydrogel structure, SEM (Merlin Gemini II, Zeiss, Germany), operating at a current of 90 pA and a voltage of 5 kV, was used for imaging the hydrogel microstructure.

### Characterizing pores in the Col-HA hydrogel structure

The microarchitecture of the Col-HA hydrogels was examined by analyzing the SEM images. The pore number and the mean pore area were quantified with the in-house Python 3.8.17 script. Data are expressed as the total number of pores within an image surface area of 32 µm^2^, shown as a mean ± standard deviation (SD), from *n* = 10 images. The surface area of the pores is presented as a median ± mean absolute deviation (MAD), from n = 10 images. Statistical significance was calculated using one-way ANOVA at α = 0.05 (**p < 0.001, ***p < 0.0001).

### Rheological properties of Col-HA hydrogels

For rheological experiments, Col-HA hydrogels were prepared as follows. Each hydrogel sample was cut from the 24-well plates using a tube with an 8 mm diameter, and the sample thickness was maintained at 2 mm. Rheological measurements were performed in oscillation mode using a parallel plate rotational rheometer (MRC302, Anton Paar, Graz, Austria). Sandblasted plates were used to avoid the hydrogels sliding during measurements. The frequency sweep measurements were conducted by applying a constant strain of γ = 1% and a frequency range of 0.1 ÷ 10 Hz. All measurements were carried out at RT with 3 repetitions.

The storage *G′* and loss *G″* moduli were obtained following the procedure published elsewhere^1,2^ and plotted as a function of frequency. To each such relationship, a double power law function was fitted^3,4^, based on which such rheological parameters like *G′, G″, tan(δ) = G″/G′,* and *ω_T_* were calculated as a mean value ± standard deviation (SD) from *n* = 9 independent hydrogel samples.

### Rheology of cancer spheroids measured by hydraulic force spectroscopy (HFS)

Rheological properties of bladder cancer spheroids were assessed using hydraulic force spectroscopy (HFS^5^). Spheroids were grown in 96-U-bottom wells in 150 µl of the corresponding culture media and collected after 3 days for HSF measurements. The number of cells/well was adjusted to obtain a spheroid diameter of 350 – 400 µm (HCV29 cells – 6000 cells/well, HT1376 cells – 1200 cells/well, and T24 cells – 10000 cells/well). For HSF measurements, an individual spheroid was captured by a glass pipette.

The HSF instrument used here was a custom-made device, which is a type of micropipette aspiration technique described elsewhere in detail^56^. Briefly, a piezoelectric micropump (TPP Ventus, USA) is connected to a water reservoir and applies a partial vacuum inside it. A second tube connects the bottom of the reservoir with the probe. Alongside it, a T-shaped junction houses an optical pressure sensor. Both the probe and the pressure sensor are connected to a fiber optic interrogator via an asymmetric splitter that delivers the light to the displacement and the pressure sensors, respectively. The light is then collected and measured by a spectrometer connected to the circulator output.

Spheroids, immersed in phosphate-buffered saline (PBS), were tested via dynamic mechanical analysis under pressure control. First samples were captured and subject to an aspiration ramp of 50 Pa/s, until a peak pressure of 300 Pa. This was followed by 20 seconds of creep, and by a series of 9 oscillatory periods between 0.05 and 20 Hz with correction for hydrodynamic drag, separated logarithmically, each of them ∼50 Pa in amplitude. In between oscillations, we added a 2-second pause. The complex modulus *E*\*, composed of storage (*E’*) and loss (*E″*) components, was computed as:

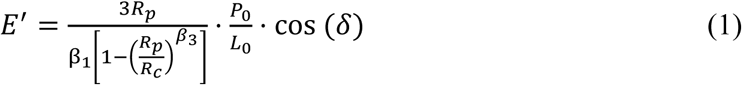

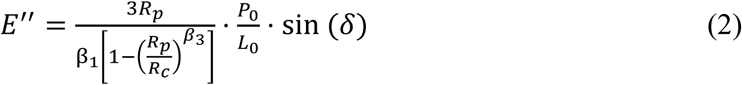

Where *R_p_* is the pipette radius, *R_c_* is the sample radius, *β_1_* is 2.0142, *β_3_* is 2.1187, *P_0_* and *L_0_* are the pressure and aspirated length amplitudes of the oscillatory periods, and *δ* their phase lag, adjusted for hydrodynamic drag^7,8^. Then, complex modulus *E*\* was converted to storage *G′* and loss *G″* moduli, using a relation *G* = *E / 2(1 + ν)*, where *ν* is a Poisson coefficient, assumed to be 0.5, for incompressible materials like cells. Each point is a mean ± SD, *n* = 10-15 spheroids. These relationships were fitted with a single-power law for *G′* and a double-power law for *G″.* Then, the data analysis was conducted as follows. Since the storage modulus *G′* displayed a weak power-law dependence across the entire frequency range, they were fitted using a single power-law model: *G′(ω) = A_0_·ω^α^,* where *ω* is the frequency of pressure oscillation. In contrast, the loss modulus *G″* exhibited a biphasic response, increasing at higher frequencies (above ∼8 Hz) and was best described by a double power-law model: 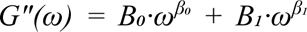. The exponent *α* provides a measure of viscoelasticity: *α* = 0 indicates a purely elastic solid, *α* = 1 a purely viscous fluid, with intermediate values reflecting gel-like behavior. At higher frequencies, the 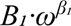 term dominates, capturing the stiffening response characteristic of semiflexible polymer networks such as collagen. The viscoelastic component was obtained by analysing *tan(δ) = G″/G′*.

### Morphology of spheroids embedded in Col-HA hydrogels

After 3 days of culture, the spheroids were collected from the U-bottom plates and placed individually on top of 1 mL of unpolymerized hydrogel solution in the individual wells of 24well plates. After polymerization (37 °C, 5% CO_2_), the samples were incubated for 24 hours in the culture medium corresponding to the cell type used. Soft pCol-HA and stiff tCol-HA hydrogels were used. For each spheroid type, 25 images of single spheroids were collected using an inverted optical microscope (Olympus IX83, Japan) with a 10× (UPL FLN2 10×/0.3) objective and equipped with a CMOS camera (Prime BSI 166 Express Scientific, Photometrics). The spheroid morphology was quantified by fitting an ellipsoid to a spheroid present on an optical image using a Python-based script. The obtained parameters were surface area, perimeter, major axis, and minor axis, which were used to calculate aspect ratio, spheroid volume, and equivalent radius.

### Calculating the energy demand to maintain a spherical shape

The obtained morphological and rhelogical data were applied to calculate the energy needed by spheroids to maintain their shape in soft and stiff Col-HA hydrogels using a Python-based script. The details on the equation used are included in the Suppl. Mat.

### Migration of cells from cancer spheroids cultured on the HCV29 monolayer

HCV29 cells were cultured in 24-well plates (TPP, Genos, Poland) for 3 days until they formed a monolayer. In parallel, spheroids formed from HCV29, T24, and HT1376 cells were grown for 3 days as described above. Afterwards, spheroids were stained alive with Cell Tracker Red (Invitrogen, 10 µM dissolved in the corresponding culture medium without FBS), for 4 hours at the CO_2_ incubator (37°C, 5% CO_2_, 95% air). Next, spheroids were placed over the HCV29 monolayer. Fluorescent images were collected after 24 h of incubation in the CO_2_ incubator.

Images were collected using an inverted optical microscope (Olympus IX83, Japan) with a 10× (UPL FLN2 10×/0.3) objective, and a U-FGW filter (λ_ex_ = 530–550 nm, λ_emi_ = 575 nm), and equipped with a CMOS camera (Prime BSI 166 Express Scientific, 01-prime-BSI-EXP, Photometrics). Next, each image of individual spheroids was analysed using a Matlab script, which fitted a sphere around the spheroid body, detected cells away from the spheroid, and calculated their distance to the edge of the spheroids.

### Cell escape from spheroids embedded in Col-HA hydrogels

Spheroids embedded in soft and stiff hydrogels were kept for 24 h, and afterwards, optical images (phase contrast) were collected using an inverted optical microscope (Olympus IX83, Japan) with a 10× (UPL FLN2 10×/0.3) objective and equipped with a CMOS camera (Prime BSI 166 Express Scientific, 01-prime-BSI-EXP, Photometrics). Next, each image of individual spheroids was analysed using a Python script, which fitted an ellipsoid around the spheroid body, detected cells away from the spheroid, and calculated their distance to the edge of the spheroid.

### Gelatin Zymography of metalloproteinase activity

Spheroids were grown in Col-HA hydrogels as described above. After 24h of culture, the supernatant was collected and used in Zymography analysis of gelatinolytic activities of metalloproteinases: MMP-2 and MMP-9. Collected supernatants were separated on a 10% SDS-PAGE gel (acrylamide-bisacrylamide; Sigma; Mini-PROTEAN, Bio-Rad, Poland) with gelatin 0.1% (1 mg/mL, Avantor Performance Materials, Poland). After running, gels were incubated in renaturing buffer (Triton X-100, 2,5%; 100 ml per gel, Avantor) with gentle incubation twice for 30 minutes at room temperature. After incubation, the gel was washed in the developing buffer composed of 50 mM Tris (pH 7.6, Merck, Poland) supplemented with 0.2M NaCl, 5 mM CaCl_2_ for 30 minutes at room temperature (RT) by gentle agitation. After washing, the developing buffer was replaced with fresh buffer and incubated at 37 °C for 48h. After incubation, the gel was stained with Coomasie Blue R-250 (0.1% Coomasie Blue R-25 (Merck, Poland), dissolved in 50% methanol (Avantor Performance Materials, Poland) and 10% acetic acid (Avantor Performance Materials, Poland)) for 2h, and washed with a solution composed of 50% methanol (Avantor Performance Materials, Poland) and 10% acetic acid (Avantor Performance Materials, Poland). Areas of protease activity appear as clear bands against a background where the protease has digested the substrate.

### Western Blot

Spheroid pellets were unfrozen, and 30 µl of a lysis buffer (composition: 1 M Tris-HCl (Merck, Poland), SDS (Merck, Poland), glycerol (Merck, Poland), and 2-mercaptoethanol (Merck, Poland)) was added. Next, they were heated for 5 min at 95-100°C in the thermoblock (Biometra, Germany) and sonicated three times for 5 s using a sonificator (power: 40%; amplitude ∼ 50 µm, 300UT, BioLogics, USA). Between sonication cycles, samples were kept on ice (4°C). The protein level in the obtained spheroid homogenates was determined using a Qubit 2.0 fluorometer (ThermoFisher Scientific, USA). A total of 40 mg of protein was loaded into each well of the electrophoretic gel. The volume of homogenate applied (in microliters) depended on its concentration, resulting in varying sample volumes across the wells. Protein separation was carried out using 4–15% Mini-PROTEAN® TGX™ Precast Protein Gels (10well format, 30 µL per well; Bio-Rad, Poland). The electrophoresis was performed in 1× Tris/Glycine/SDS buffer prepared from a 10×concentrate (Bio-Rad, Poland) under the following conditions: 120 V for approximately 1.5 hours at +4°C. For the Western blot transfer, the transfer buffer was prepared per liter using 700 mL of water, 100 mL of 10× Tris/Glycine buffer (Bio-Rad, Poland), and 200 mL of methanol (Avantor Performance Materials, Poland). The transfer to PVDF membrane (Roche, Switzerland) was conducted at a current intensity of 0.15 mA for approximately 20 hours at +4°C. The following antibodies were applied: mouse monoclonal antibody against vinculin (hVIN-1, Merck, Poland), monoclonal antibody against E-cadherin (DECMA-1, Invitrogen, USA), mouse monoclonal antibody against N-cadherin (Merck, Poland), and rabbit monoclonal antibody against syndecan-4 (Cell Signalling Technology, USA). As a control, the level of GAPDH was assessed by using a rabbit monoclonal antibody (Cell Signaling Technology, USA). Bands were visualized using alkaline phosphatase-conjugated anti-rabbit or anti-mouse secondary antibodies (Merck, Poland) and detected by a colorimetric reaction with nitro blue tetrazolium (NBT)/5-bromo-4-chloro-3indolyl phosphate (BCIP) substrate (Roche, Switzerland). To detect the presence of collagen I and IV in the studied bladder cancer cells, cell homogenates were prepared in an analogous way to spheroid homogenates. Antibodies against: collagen I (Invitrogen, USA) diluted 1:500, collagen IV (Merck, Poland) diluted 1:100. The presence of primary antibody was revealed with horseradish peroxidase-conjugated secondary antibody diluted 1:20000 (Cell Signalling Technology, USA) and visualized with an Enhanced Chemiluminescence (ECL) detection system (Bio-Rad, Poland).

### Fluorescence imaging of Col-HA hydrogels with spheroids

The volume of 160 μL of Col-HA hydrogel solution was transferred into 24-well glass-bottom microplates (Biokom, Poland). Individual spheroids were placed on top of an unpolymerized solution and left for polymerization (37 °C, 5% CO_2_, 60 min). Samples were incubated for 24h, 48h, and 72 h in the culture medium corresponding to the cell type. After each time point, the culture medium was removed, followed by rinsing three times in PBS. Samples were fixed with a 3.7% paraformaldehyde solution (Merck, Poland) in PBS for 20 min, and then rinsed 3 times in PBS. Next, the samples were incubated with 1 μg/mL mouse monoclonal antibody against collagen I (5D8-G9, Invitrogen, USA) for 6 h at 25 °C, followed by five rinses in PBS for 30 minutes each. They were incubated with 2 μg/mL Alexa Fluor 555 goat anti-mouse IgG secondary antibody (Invitrogen, USA) for 6 h at 25 °C and again rinsed 5 times with PBS. Cell nuclei in spheroids were stained with a 1 μg/mL DAPI solution (BD Bioscience, USA) for 30 minutes. After rinsing with PBS, the samples were imaged immediately. Fluorescence images (2048 px × 2048 px) were acquired using an inverted microscope Olympus IX83 (Olympus, Japan) with a 40× objective (LUC PLAN FLN 40×/0.60), and a CMOS camera type Prime BSI 166 Express Scientific (01-prime-BSI-EXP, Photometrics). A U-FGW filter (*λ_ex_* = 530–550 nm, *λ_emi_* = 575 nm) was utilized to image collagen fibers (Alexa Fluor 555), whereas a U-FUW filter (*λ_ex_* = 340–390 nm, *λ_emi_* = 420 nm) was applied to record images of stained cell nuclei (Hoechst 34580).

### Real-time quantitative polymerase chain reaction (RT-qPCR)

The total RNA of spheroid pellets, stored at -80°C until RNA isolation, was isolated using Maxwell RSC RNA simplyRNA Cells Kit (Promega, Poland). After concentration and purity determination with the use of Nanodrop (ThermoFisher Scientific, USA), RNA was reverse transcribed to cDNA using MultiScribe Reverse Transcriptase (High-Capacity cDNA Reverse Transcriptase Kit, ThermoFisher Scientific, USA). For gene expression analysis, we used qPCR with validated TaqMan assays (ThermoFisher Scientific, USA) using the hydrolysis probe method. We analysed two transcripts for genes: SDC4 (id: Hs01120908_m1) and HK2 (id: Hs00606086_m1) in three groups of samples: spheroids alone, spheroids embedded in soft Col-HA hydrogels, and spheroids embedded in stiff Col-HA hydrogels. As an endogenous control, we used 18S ribosomal RNA (id: Hs03003631_m1). Briefly, 10 times diluted cDNA was added to 1X TaqMan gene expression Master Mix (ThermoFisher Scientific, USA) with a specific set of primers used to detect the SDC4, HK2 transcripts, or the 18S endogenous control. Real-time reaction was carried out on CFX384 thermal cycler (BioRad, Poland) for 40 cycles at standard thermal protocol, processed in triplicate with endogenous control on the same 384 format plate. The Cq data (cycle threshold set up at 500 RFU) were analysed using R in the RStudio environment. Gene expression was measured as a relative expression using the ddCt method with 18S rRNA as an internal control (reference gene) and the spheroid alone group as a reference group^7^. Main calculations were performed using the“ pcr” package^8^ with the control group (spheroid alone group) set as 1. The data visualization was obtained by using OriginPro 2022.

## Suppl. Note S1. Viability of cells in spheroids

Cell viability was determined using a commercially available Live/Dead Cell Double Staining Kit (Merck, Poland). After 24 h of spheroid incubation in the hydrogels, the culture medium was removed from the samples, followed by careful rinsing with phosphate-buffered saline (PBS). Next, 10 µL of calcein acetoxymethyl ester (Calcein-AM) and 0.5 µL propidium iodide (PI) were mixed with 5 mL of PBS. Next, 300 µL of the prepared assay was added to each sample, i.e., Col-HA hydrogel with embedded spheroids, prepared as described in *Materials & Methods*. Samples were then incubated for 15 min and imaged immediately afterward with an epi-fluorescent microscope (Olympus IX83, Japan) with a UPL FLN2 10×/0.3 objective, and equipped with a CMOS camera (Prime BSI 166 Express Scientific, Photometrics). A U-FBW filter (λ_ex_ = 460–495 nm, λ_emi_ = 510 nm) was used to detect live cells (Calcein-AM), while a UFGW filter (λ_ex_ = 530–550 nm, λ_emi_ = 575 nm) was used to identify dead cells (PI). The cell viability was repeated 3 times.

**Suppl. Fig. S1.**
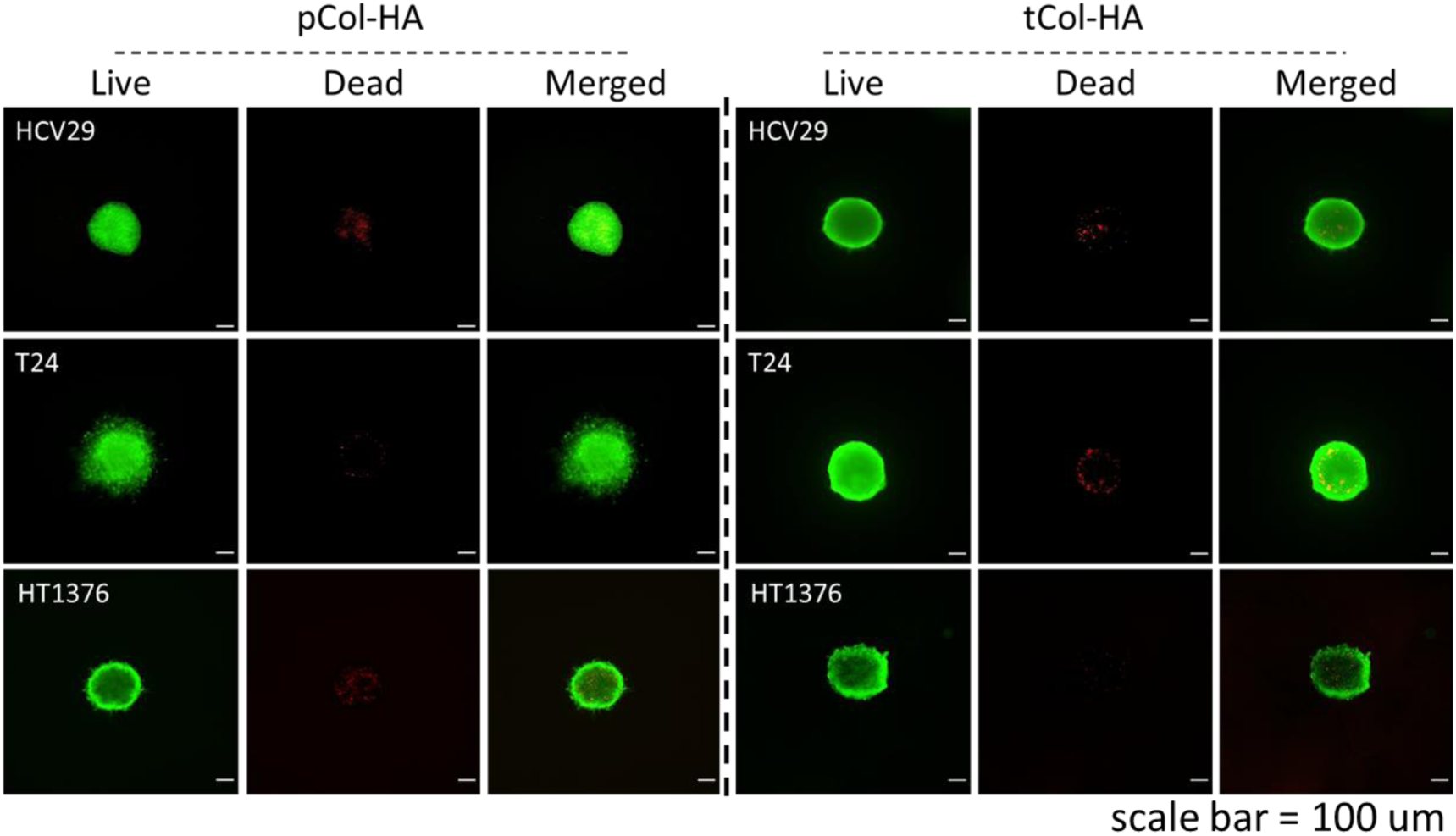
Viability of cells inside spheroids embedded in soft (pCol-HA) and stiff (tCol-HA) hydrogels, assessed as described in **Suppl. Note S1**.

## Suppl. Note S2. Rheology of Col-HA hydrogels with embedded spheroids

For rheological experiments, Col-HA hydrogels with embedded 48 spheroids of each cell type were prepared as follows. Each hydrogel sample was cut from the 24-well plates using a tube with an 8 mm diameter, and the sample thickness was maintained at 2 mm. Rheological measurements were performed in oscillation mode using a parallel plate rotational rheometer (MRC302, Anton Paar, Graz, Austria). Sandblasted plates were used to avoid the hydrogels sliding during measurements. The frequency sweep measurements were conducted by applying a constant strain of γ = 1% and a frequency range of 0.1 ÷ 10 Hz. All measurements were carried out at RT with 3 repetitions.

**4. Suppl. Fig. S2.**
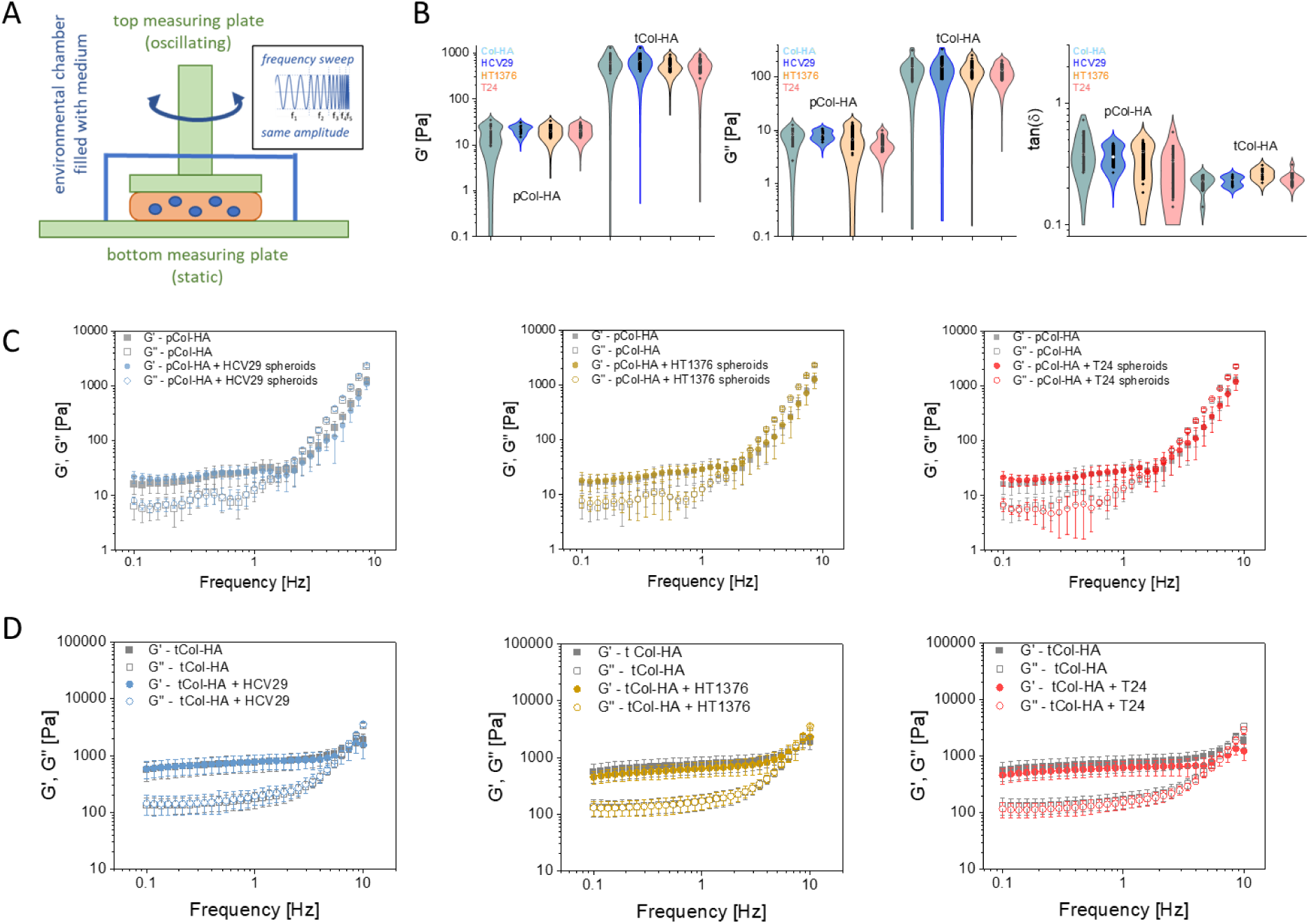
(A) The idea of frequency-sweep rheology for samples composed of spheroids embedded in Col-HA hydrogels, measured using a rheometer (Anton Paar, MRC 302). (B) Calculated rheological parameters, i.e*., G*’, *G″*, and *tan(δ)*, obtained for soft pCol-HA and stiff tCol-HA hydrogels (n = 9 hydrogels per group, 48 spheroids per hydrogel). (C) The resulting storage (*G′*) and loss (*G*″) moduli are plotted as a function of frequency of applied oscillations. Each point represents a mean±SD (n=9 hydrogels per type).

**Supplementary Table S1.** Morphological parameters calculated for spheroids embedded in soft and stiff Col-HA hydrogels for 24h (data are presented as a mean ± standard deviation for n = 25 spheroids). These include perimeter, lengths of the major and minor axes, aspect ratio (minor/major axis), equivalent radius (defined as *√(major × minor axis)*), and an estimated spheroid volume, calculated as the volume of a sphere with the equivalent radius.

| *Spheroids* | *Surface area*<br>[$\mu\text{m}^2$] | *Perimeter*<br>[$\mu\text{m}$] | *Major axis*<br>[$\mu\text{m}$] | *Minor axis*<br>[$\mu\text{m}$] | *Equivalent radius*<br>[$\mu\text{m}$] | *Aspect ratio* |
| --- | --- | --- | --- | --- | --- | --- |
| pCol-HA |  |  |  |  |  |  |
| HCV29 | 186 206 ± 48192 | 4477 ± 1315 | 512 ± 68 | 469 ± 60 | 490 ± 62 | 1.09 ± 0.07 |
| HT1376 | 330 636 ± 131049 | 6078 ± 3765 | 666 ± 145 | 613 ± 141 | 639 ± 142 | 1.09 ± 0.06 |
| T24 | 485 740 ± 12496 | 8788 ± 3330 | 850 ± 274 | 774 ± 248 | 810 ± 260 | 1.08 ± 0.04 |
| tCol-HA |  |  |  |  |  |  |
| HCV29 | 70 280 ± 58100 | 1497 ± 863 | 309 ± 119 | 253 ± 110 | 279 ± 114 | 1.24 ± 0.14 |
| HT1376 | 110 301 ± 3156 | 3352 ± 3024 | 373 ± 164 | 327 ± 151 | 349 ± 157 | 1.15 ± 0.11 |
| T24 | 106 160 ± 79091 | 4664 ± 2547 | 503 ± 135 | 435 ± 116 | 497 ± 124 | 1.16 ± 0.10 |

**Supplementary Table S2.**
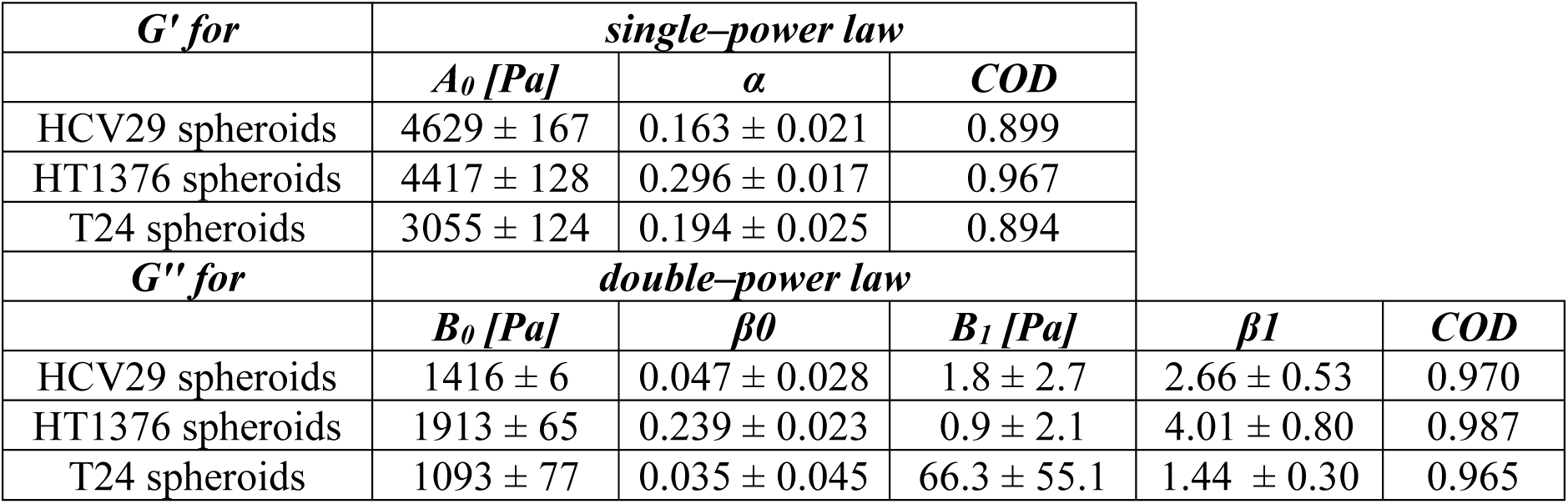
Table 1. Power-law parameters derived from hydraulic force spectroscopy of bladder cancer spheroids in PBS. Values are presented as fitted parameter±fit error; COD denotes the coefficient of determination (R²). Single and double power laws describe the viscoelastic response of soft materials as a function of angular frequency (*ω*)^5–9^.

| *G' for* | *single-power law* |  |  |  |  |
| --- | --- | --- | --- | --- | --- |
|  | *A<sub>0</sub> [Pa]* | *$\alpha$* | *COD* |  |  |
| HCV29 spheroids | 4629 ± 167 | 0.163 ± 0.021 | 0.899 | | |
| HT1376 spheroids | 4417 ± 128 | 0.296 ± 0.017 | 0.967 | | |
| T24 spheroids | 3055 ± 124 | 0.194 ± 0.025 | 0.894 | | |
| *G'' for* | *double-power law* |  |  |  |  |
|  | *B<sub>0</sub> [Pa]* | *$\beta_0$* | *B<sub>1</sub> [Pa]* | *$\beta_1$* | *COD* |
| HCV29 spheroids | 1416 ± 6 | 0.047 ± 0.028 | 1.8 ± 2.7 | 2.66 ± 0.53 | 0.970 |
| HT1376 spheroids | 1913 ± 65 | 0.239 ± 0.023 | 0.9 ± 2.1 | 4.01 ± 0.80 | 0.987 |
| T24 spheroids | 1093 ± 77 | 0.035 ± 0.045 | 66.3 ± 55.1 | 1.44 ± 0.30 | 0.965 |

## Suppl. Note S3. Model of energetic costs required to maintain the spheroid shape in the mechanically different microenvironments

To calculate how much energy spheroids require to maintain a shape, being embedded in a viscoelastic environment, we developed a simplified model of a viscoelastic sphere (radius *R_S_*, volume *V_S_,* storage modulus *G’_S_*, loss modulus *G″_S_*) embedded within an infinite thick viscoelastic matrix (storage modulus *G’_M_*, loss modulus *G″_M_)*. We consider steady-state conditions in which the sphere remains stationary within a viscoelastic matrix. The only force acting on the sphere is a compressive shear strain of scalar amplitude (*ε\**), describing the extent to which the sphere is compressed. The sphere and the matrix are coupled, so the deformation of the sphere does not equal the external strain. Part of the deformation is transferred to the matrix, while the sphere itself absorbs the remainder (*ε_S_)*.

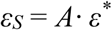

Here, *A* characterizes the relationship between the deformations of the matrix and the sphere. Physically, *A* arises from the balance of shear forces between the inclusion and the surrounding matrix. One can imagine the sphere as a spring with elastic modulus *E_S_* and the matrix as a spring with modulus *E_M_* (*E = 2·G·(1 + ν)*, where *ν* is the Poisson ratio), such that the applied deformation is shared between them in proportion to their respective mechanics. According to Eshelby’s theory^10^, for a perfectly spherical inclusion embedded in an infinite matrix, a simple expression for the shear deformation is:

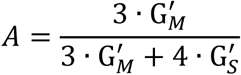

In the case of a very soft sphere (i.e., *G’_S_ << G’_M_*), *A* = 1, which means that the surrounding matrix deforms the sphere. When *A* = 0, a sphere is practically not deforming.

The elastic energy, gathered in a sphere, is:

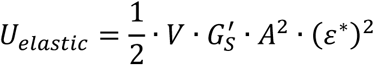

In the case of the sphere expansion, volumetric strain is rather observed than pure shear; therefore, the analysis must be extended to include viscoelastic effects by introducing complex shear and bulk moduli, *G\** and *K\**, to account for additional contribution from volumetric deformation *A_vol_*, which is:

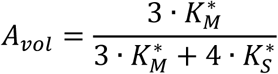

where:

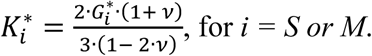

Thus, two energy components can be obtained:

Energy stored in the sphere: 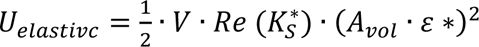

Energy dissipated due to viscosity: 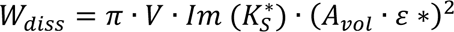

## Suppl. Note S4. Cell line characteristics

Non-malignant HCV29 cells, which were originally derived from a non-malignant part of the irradiated, cancer-free bladder transitional epithelium (urothelium) of the patient suffering from bladder carcinoma^11^, retain some features of normal cells. These include epithelial-like morphology, adherent growth forming monolayers, and well-differentiated cytoskeleton populated with actin stress fibers spanning the whole cell body^12^. They are more rigid than cancer cells at both cellular and spheroid levels^13,14^, displaying more solid-like behavior^14^. We also show in previous studies that they express N-cadherins (large expression^15^), E-cadherins (low expression^15^), and SDC-4 (low expression^16^).

HT1376 cells were obtained from a transurethral resection of invasive (grade 3) transitional cell carcinoma of a 58-year-old Caucasian women^17^. These cells, characterized by poorly differentiated actin filaments without the presence of actin stress fibers, show the largest deformability as compared to HCV29 and T24 cells^13,14^, displaying more fluid-like behavior^14^. They also express a large level of E-cadherins and no N-cadherins, and the highest level of SDC-4^16^. Their migratory activity was the lowest among the studied bladder cancer cells.

T24 cells were derived from a high-grade, invasive transitional cell carcinoma (TCC)^18^. These cells are characterized by moderately differentiated actin filaments with fewer abundant stress fibers than in the case of HCV29 cells, displaying deformability between HCV29 and HT1376 cells^13,14^. They express a large level of N-cadherins^15^, no E-cadherins^15^, and the level of SDC4 between HCV29 and HT1376 cells. Their migratory activity was the highest among the studied bladder cancer cells.

**Suppl. Fig. S3.**
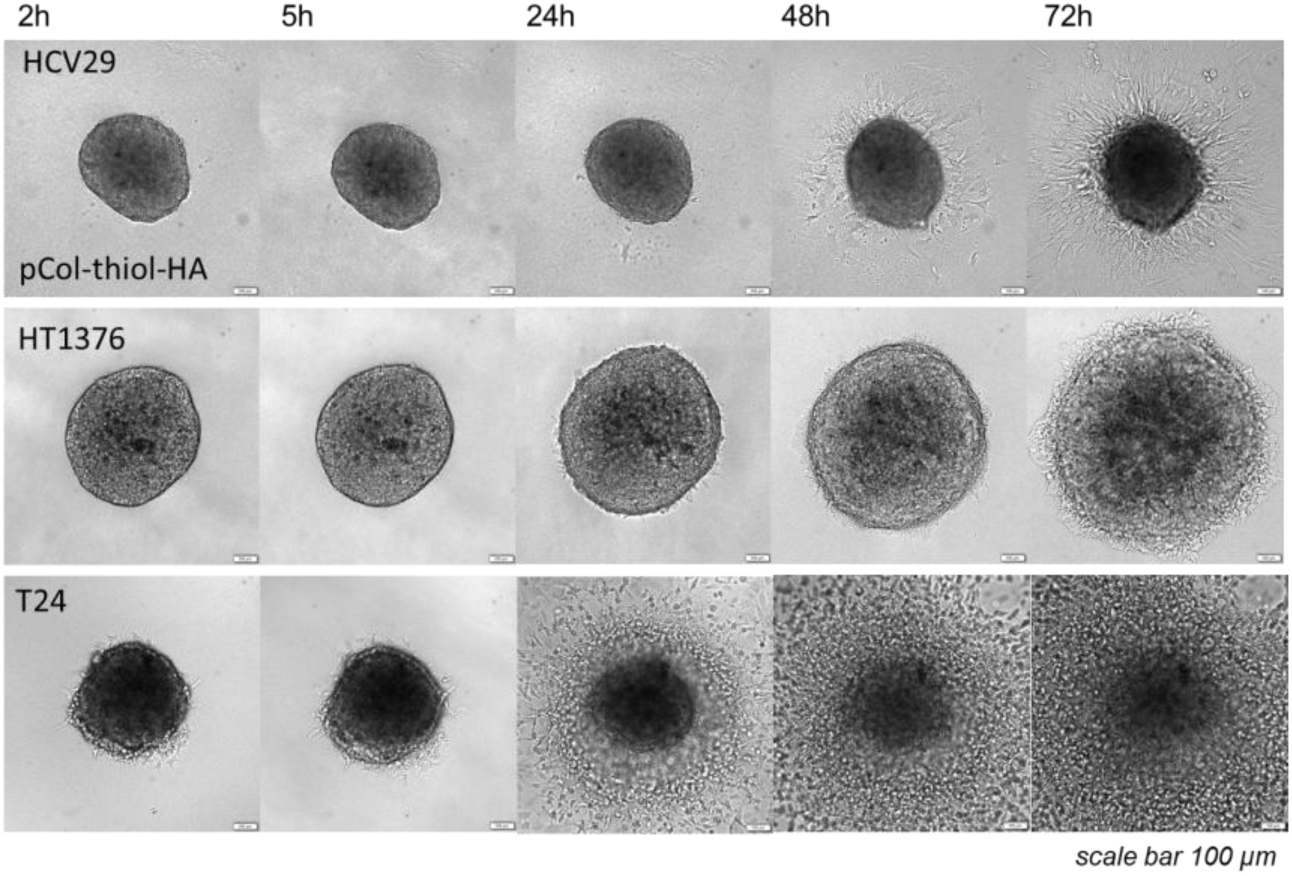
Exemplary time-dependent expansion of bladder cancer spheroids in soft pCol-HA hydrogels – the same spheroids were traced up to 72h.

**Suppl. Fig. S4.**
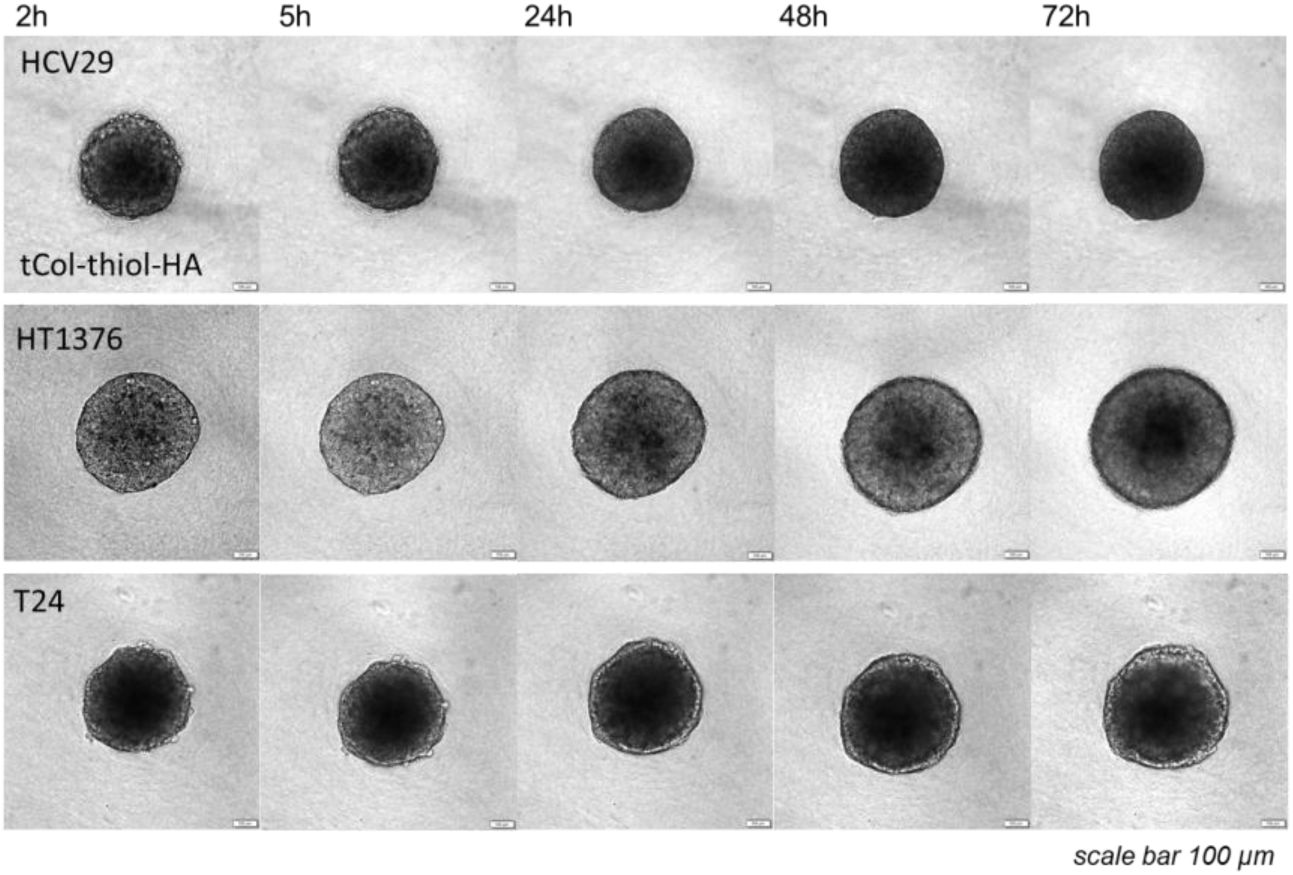
Time-dependent expansion of bladder cancer spheroids in stiff tCol-HA hydrogels – the same spheroids were traced up to 72h.

**Suppl. Fig. S5.**
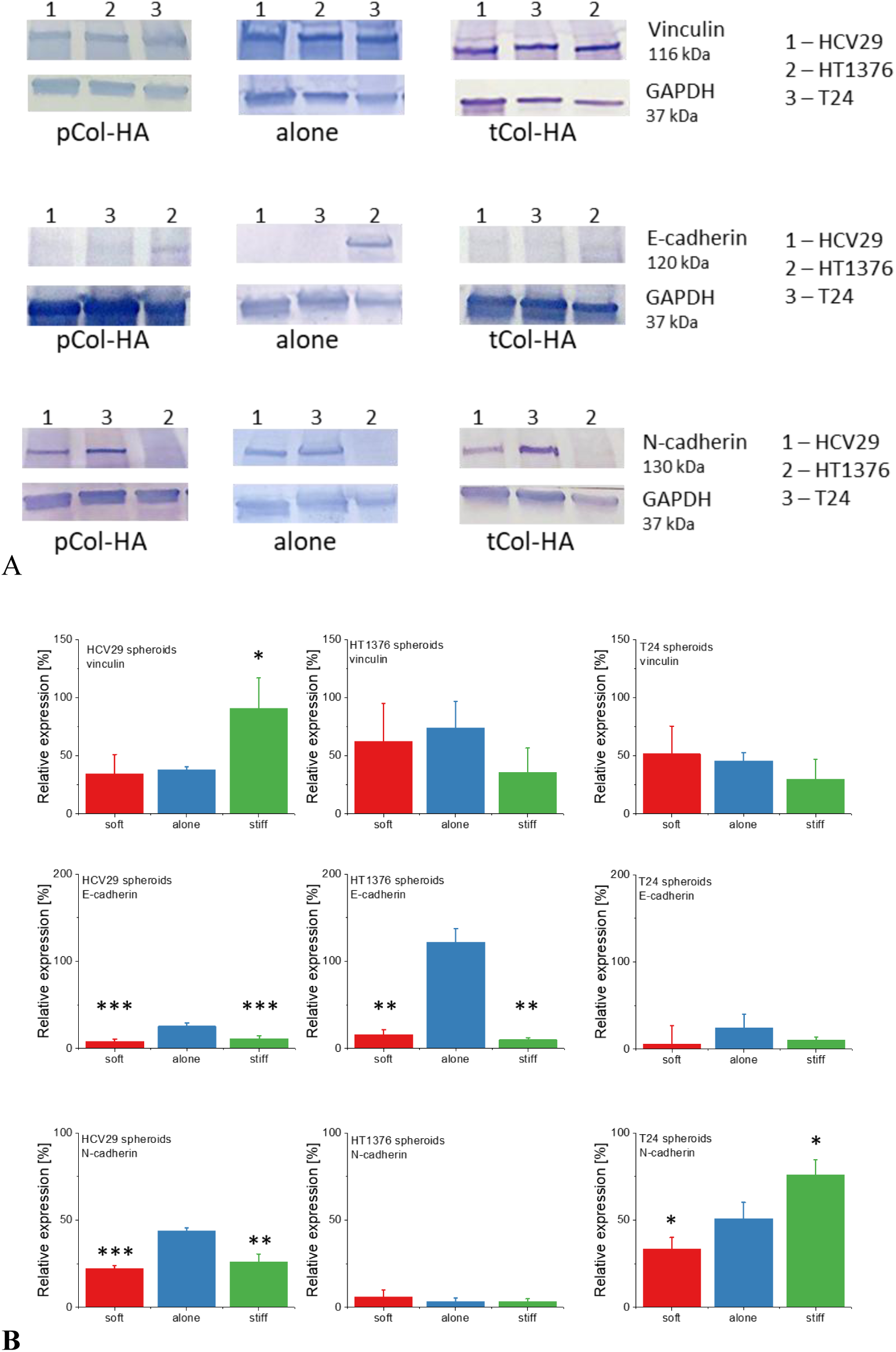
A) Exemplary bands from Western blots, presented as a studied protein together with the corresponding control (GAPDH). B) Relative expression of vinculin, E-cadherin, and N-cadherin at the protein level, calculated based on densitometry (Image J). Three groups were studied: spheroids alone, spheroids embedded in the soft Col-HA hydrogels, and spheroids embedded in the stiff Col-HA hydrogels (n = 3 repetitions). The results are presented in relation to the expression in spheroids grown only in culture medium (spheroids alone group, i.e., control, to which expression levels were normalized). Statistical significance was obtained from an unpaired Student t-test at the level of 0.05.

**Suppl. Fig. S6.**
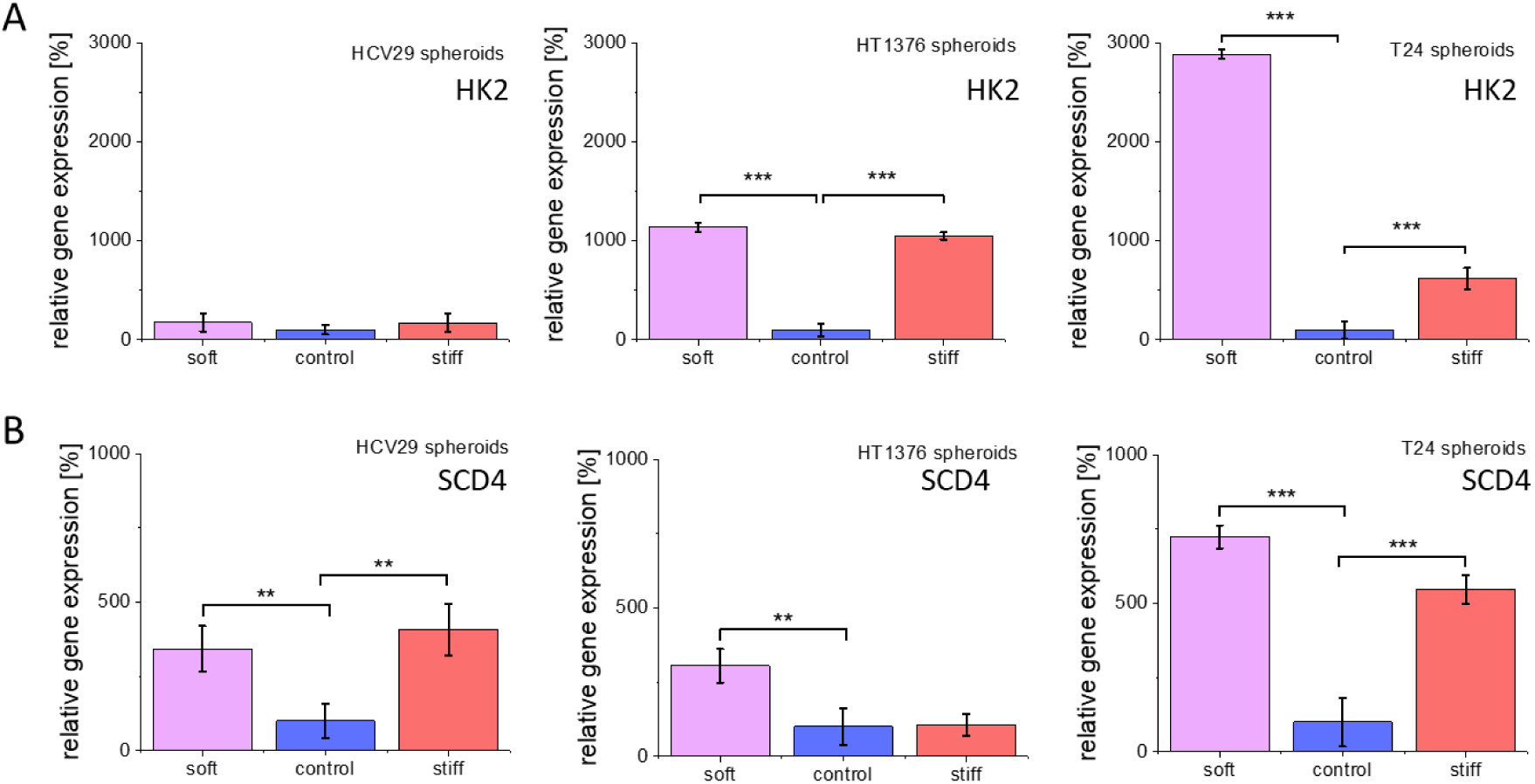
Relative expression of HK2 and SDC4 at the mRNA level, calculated as described in *Materials and Methods*. Three groups were studied: spheroids alone, spheroids embedded in the soft Col-HA hydrogels, and spheroids embedded in the stiff Col-HA hydrogels. The results are presented in relation to the expression in spheroids grown only in culture medium (spheroids alone group, i.e., control, to which expression levels were normalized). Statistical significance was obtained from an unpaired Student t-test at the level of 0.05.

